# Molecular insights into differentiated ligand recognition of the human parathyroid hormone receptor 2

**DOI:** 10.1101/2021.06.09.447485

**Authors:** Xi Wang, Xi Cheng, Lihua Zhao, Yuzhe Wang, Chenyu Ye, Xinyu Zou, Antao Dai, Zhaotong Cong, Jian Chen, Qingtong Zhou, Tian Xia, Hualiang Jiang, Eric H. Xu, Dehua Yang, Ming-Wei Wang

## Abstract

The parathyroid hormone receptor 2 (PTH2R) is a class B1 G protein-coupled receptor (GPCR) involved in regulation of calcium transport, nociception mediation, and wound healing. Naturally occurring mutations in PTH2R were reported to cause hereditary diseases, including syndromic short stature. Here we report the cryo-electron microscopy structure of PTH2R bound to its endogenous ligand, tuberoinfundibular peptide (TIP39), and a heterotrimeric G_s_ protein at a global resolution of 2.8 Å. The structure reveals that TIP39 adopts a unique loop conformation at N terminus and deeply inserts into the orthosteric ligand-binding pocket in the transmembrane (TM) domain. Molecular dynamics (MD) simulation and site-directed mutagenesis studies uncover the basis of ligand specificity relative to three PTH2R agonists, TIP39, PTH, and PTH-related peptide (PTHrP). We also compare the action of TIP39 with an antagonist lacking six residues from the peptide N terminus, TIP(7–39), which underscores the indispensable role of the N terminus of TIP39 in PTH2R activation. Additionally, we unveil that a disease-associated mutation G258D significantly diminished cAMP accumulation induced by TIP39. Together, these results not only provide structural insights into ligand specificity and receptor activation of class B1 GPCRs, but also offer a foundation to systematically rationalize the available pharmacological data to develop novel therapies for various disorders associated with PTH2R.

## Introduction

Class B1 G protein-coupled receptors (GPCRs) comprise 15 members involved in a wide spectrum of physiological functions (1, 2). A number of them are validated drug targets for different human diseases, such as osteoporosis, type 2 diabetes, obesity, psychiatric disorders, and migraine. Among them are two types of parathyroid hormone (PTH) receptors (PTH1R and PTH2R), whose action are mediated by coupling primarily to the stimulatory G protein (G_s_) (3, 4). Expressed in the central and peripheral nervous systems, PTH2R is a key mediator of nociception, wound healing and maternal behavior (5–8). In addition, recent studies have shown that it regulates calcium transport and influences keratinocyte differentiation, pointing to its potential in the treatment of Darier disease or Hailey-Hailey disease (9). In addition, naturally occurring PTH2R mutations have been linked to familial early-onset generalized osteoarthritis, syndromic intellectual disability and syndromic short stature (10, 11). The latter is presently being treated with recombinant human growth hormone (12).

PTH receptors have three endogenous ligands, namely, tuberoinfundibular peptide of 39 residues (TIP39), PTH and parathyroid hormone-related peptide (PTHrP). Unlike PTH and PTHrP that mainly expressed in peripheral systems, TIP39-containing neuronal cell bodies have been identified in the subparafascicular area and the medial paralemniscal nucleus (13). Both PTH and PTHrP are implicated in skeletal development, calcium homeostasis and bone turnover (14). In fact, PTH(1–34) and abaloparatide, a variant of PTHrP(1–34) (15), are FDA approved drugs for osteoporosis. Discovered in the bovine hypothalamus, TIP39 contains two identical and several similar residues common to PTH and PTHrP. However, there is no evidence to suggest that TIP39 plays a role in mineral or bone metabolism. In contrast to PTH that indistinguishably activates both receptors, TIP39 is selective for PTH2R (13, 16), while PTHrP only has a weak action on PTH2R (3, 4, 13). Deletion of six residues from the N terminus of TIP39 results in a PTH2R antagonist, TIP(7–39) (17). However, the underlying mechanism by which PTH2R selectively recognizes these related but distinct peptides is largely unknown. Although newly solved cryo-electron microscopy (cryo-EM) structure of LA-PTH-PTH1R-G_s_ complex offers valuable insights into PTH recognition and receptor activation (18), questions remain relative to their applicability to PTH2R. Thus, we determined the single-particle cryo-EM structure of the human PTH2R in complex with TIP39 and a heterotrimeric G_s_ protein at a global resolution of 2.8 Å. Together with molecular dynamics (MD) simulation results, it provides an in-depth understanding of the structural basis of ligand specificity and PTH2R activation.

## Results

### Overall structure

As shown in Fig. 1 and Figs. S1-S2, the final model of the PTH2R-G_s_ complex contains the first 34 amino acids of TIP39, the PTH2R (residues from Thr31^ECD^-Ser434^8.64b^) (class B1 GPCR numbering in superscript) (19), a dominant-negative human Gα_s_ including eight mutations (S54N, G226A, E268A, N271K, K274D, R280K, T284D and I285T), except for the α-helical domain (AHD), rat Gβ1, bovine Gγ2 and nanobody Nb35. Excluding the extracellular domain (ECD), the side chains of a majority of residues were well defined in the EM density maps (Fig. 1A, Fig. S3 and Table S1). The overall structure of this complex is similar to that of other activated class B1 GPCRs such as LA-PTH– PTH1R–G_s_ (18), GLP-1–GLP-1R–G_s_ (20) and glucagon–GCGR–G_s_ (21) with Cα root mean square deviation (RMSD) values of 0.98 Å, 0.72 Å, and 0.94 Å for the whole complex, respectively.

**Figure 1.**
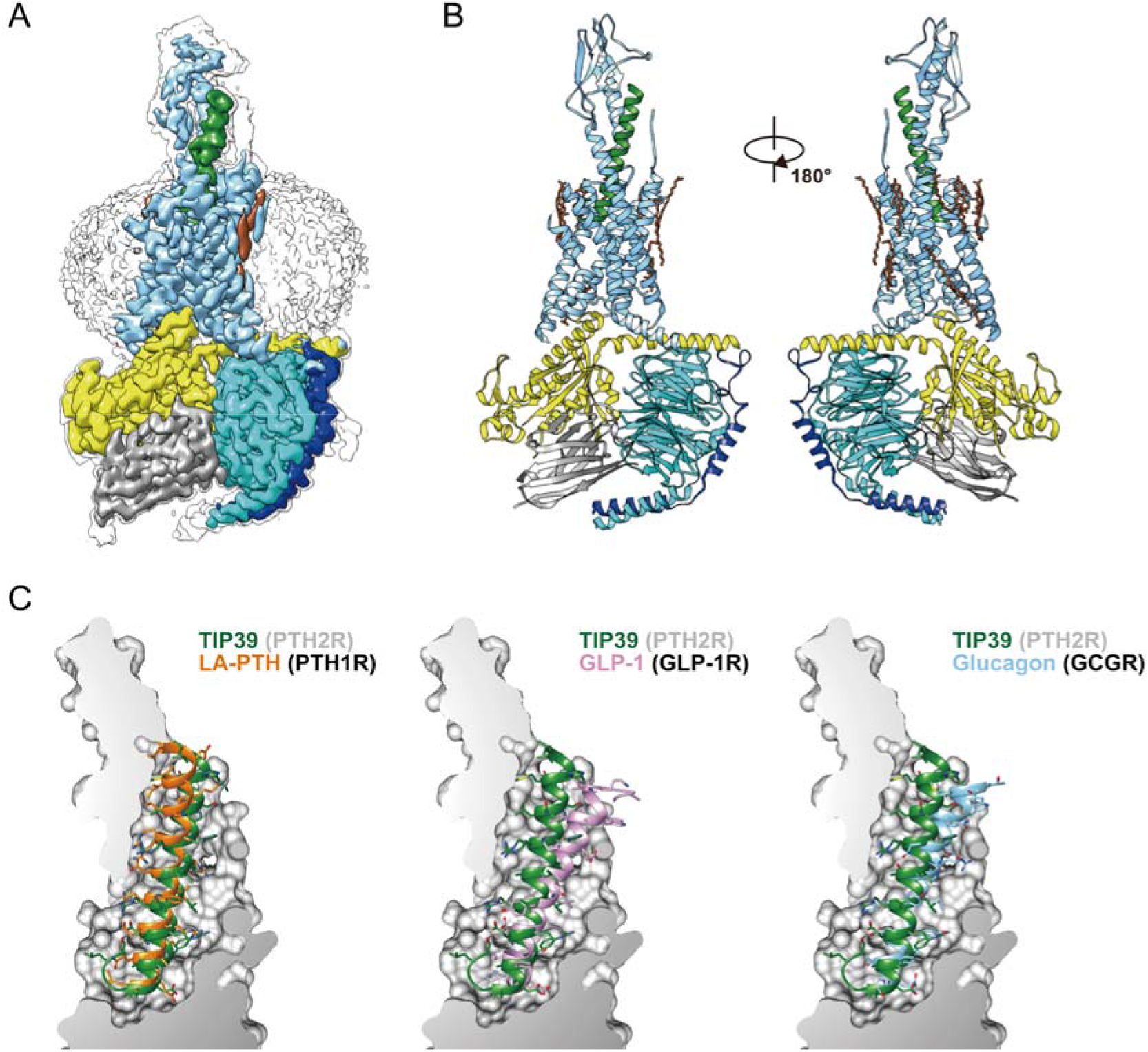
The overall cryo-EM structure of the TIP39–PTH2R-G_s_ complex. **(*A*)** Cut-through view of the cryo-EM density map that illustrates the TIP39-PTH2R-G_s_ complex and the disc-shaped micelle. The unsharpened cryo-EM density map at the 0.06 threshold shown as gray surface indicates a micelle diameter of 11 nm. The colored cryo-EM density map is shown at 0.12 threshold. **(*B*)** Model of the complex as a cartoon, with TIP39 as helix in green. The receptor is shown in blue, Gα_s_ in yellow, Gβ subunit in cyan, Gγ subunit in navy blue and Nb35 in gray. **(*C*)** The binding pocket of PTH2R accommodates peptide ligands of class B1 receptors. TIP39 is compared with LA-PTH (left), GLP-1 (middle) and Glucagon (right), respectively.

A notable structural difference occurs in the TMD ligand-binding pocket of these receptors. Fig. 1C and Fig. S4 illustrate the shapes and the sizes of the TMD pockets and their cognate ligands. The interfacing structure of TIP39-PTH2R buried areas is 2,068 Å^2^, 65% of which was contributed by the N-terminal half of TIP39. Different from a typical peptide in the class B1 GPCR subfamily that adopts an extended helix with its N-terminus inserted deeply into the TMD, TIP39 exhibits a single amphipathic α-helix from Leu4^P^ (P indicates that the residue belongs to peptide) to Leu34^P^, with Leu4^P^ being the deepest residue within the receptor core, and adopts a closed loop at the peptide N-terminus (the first three residues) surrounded by TM5, TM6, ECL2 and ECL3 (Fig. 2A). In addition, unlike other class B1 GPCRs, PTH2R has an extended TM1 helix capable of interacting with a peptidic ligand. Diverse ECD positions in PTH2R and other class B1 GPCRs also presumably adjust individual peptide helix to respective TMD pocket in a manner that is specific for each receptor (Fig. 1C, Fig. S4). In contrast to the ECL1 of growth hormone releasing hormone receptor (GHRHR) that stretches around GHRH to form broad interactions, no structural features in the ECL1 region of PTH2R were observed. This subtle difference supports our previous hypothesis that different activation requirement exists in class B1 GPCRs (22).

**Figure 2.**
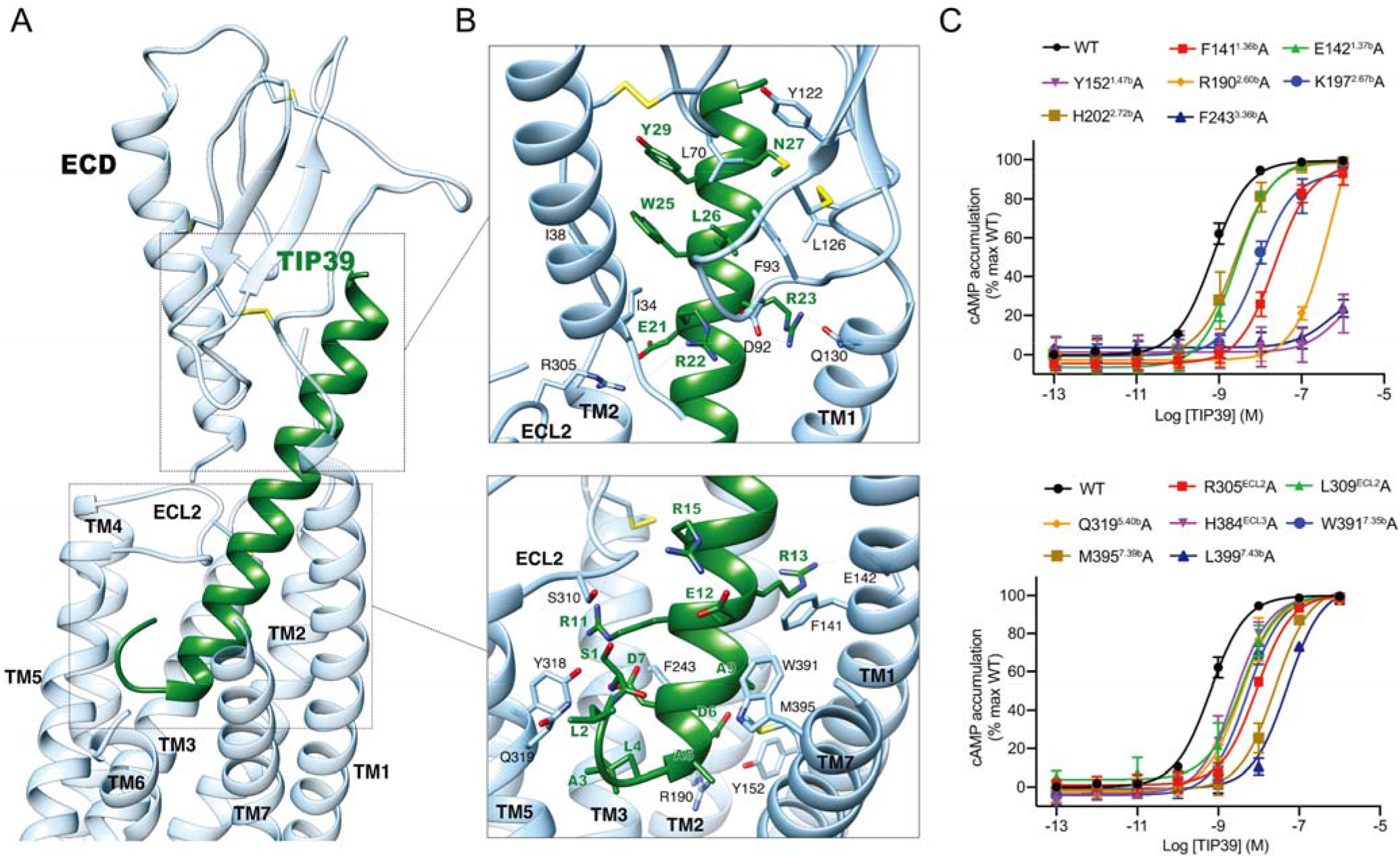
Molecular recognition and ligand specificity of PTH2R. **(*A*)** Overall contacts between PTH2R (blue) and TIP39 (green). **(*B*)** Detail contacts between PTH2R (blue) and TIP39 (green) within the ECD or the TMD. Key residues are shown as sticks. **(*C*)** Effects of receptor mutations on TIP39-induced cAMP accumulation. Data shown are means ± S.E.M. of at least three independent experiments.

With respect to the PTH2R-G_s_ interface, the outward movement of TM6 leads to a large opening of the cytoplasmic cavity for G_s_ coupling. The overall assembly of receptor-G_s_ complexes is very similar among class B1 GPCR structures solved to date (18, 23–25). In this study, the PTH2R-G_s_ complex is anchored with the α5-helix of Gα_s_, which fits snugly into the cytoplasmic cavity of the TMD. Additional contacts are observed between the extended helix 8 and the Gβ subunit (Fig. S5). A number of detailed side chain interactions are visible in the receptor-G_s_ interface (Fig. S5). The side chain of Glu392 and the last helical residue of the α5 helix in G°_s_ form a capping interaction with backbone amine of helix 8. Arg385 and Asp381 at the middle of the α5 helix in Gα_s_ make charged interactions with Glu346^ICL3^ and Lys343^5.64b^ of TM5, respectively. The carboxylate group at the C-terminal end of the α5 helix in Gα_s_ forms a salt bridge with Lys360^6.37b^ of TM6. Besides these polar and charged interactions, hydrophobic residues Leu388, Tyr391, Leu393 and Leu394 pack tightly against the hydrophobic surface comprised of residues of TM2, TM3, TM5, TM6 and TM7. Additionally, like other class B1 GPCRs, the α5 helix of G°s also interacts with ICL2 and helix 8 of PTH2R (Fig. S5).

### Ligand specificity

An extensive network of complementary polar and non-polar contacts between TIP39 and PTH2R was observed (Fig. 2 and Table S2). Pointing to the receptor core, Ser1^P^ forms one hydrogen bond with ECL2 (Ser310^ECL2^) via its side chain, and has its amine terminus interact with the α-helix part (Asp7^P^) of TIP39. Asp6, a highly conserved residue in glucagon-like peptides (26), makes one hydrogen bond and a salt bridge with Tyr152^1.47b^ and Arg190^2.60b^, respectively, in line with abolished or decreased potencies for TIP39 observed in mutants Y152A and R190A (by 794-fold) (Fig. 2C, Tables S3 and S4). Meanwhile, Glu21^P^ forms a salt bridge with Arg305^ECL2^, consistent with a 16-fold reduction of TIP39 potency in mutant R305A (Fig. 2C, Tables S3 and S4). Non-polar interactions between TIP39 and PTH2R TMD are mainly contributed by the extracellular portions of TMs 1, 2 and 7, involving Phe141^1.36b^, Lys197^2.67b^, Phe243^3.36b^, Met395^7.39b^ and Leu399^7.43b^. Removal of the non-polar contacts by alanine substitutions lowered the peptide potency by 12 ∼ 80-fold (Fig. 2C and Tables S2-S4). Of interest, the TM1 of PTH2R bends down towards TIP39, resulting in polar interactions between Arg23^P^ and Gln130^1.25b^, and shifting the peptide C-terminal region towards ECL1, while the ECD clasps this region (residues 22 to 39) with massive hydrophobic contacts and several polar interactions (Fig. 2B and Table S2).

Structural comparison of TIP39–PTH2R–G_s_ and LA-PTH–PTH1R–G_s_ complexes reveals distinct features of the ligand recognition pattern between PTH1R and PTH2R. To specifically accommodate TIP39, PTH2R reforms its peptide-binding pocket by reorganizing the conformations of ECL3 and the extracellular parts of TM1 and TM7, as well as adopts receptor-specific amino acids at multiple positions that directly interact with the peptide. ECL3 is unstructured in PTH2R but is well solved in PTH1R that forms several additional direct contacts with the N-terminal portion of the bound LA-PTH (Figs. 2A and 3A). Such differences might contribute to the greater mobility of ECL3 in PTH2R that moves outward in response to the unique loop conformation at the N terminus of TIP39. Consequently, the extracellular tip of TM7 in PTH2R also shifts outward by 2 Å (measured at the Cα of Trp^7.35b^) thereby decreasing the contacts with the bound TIP39. Meanwhile, the extracellular tip of TM1 in PTH2R is extended by six residues, allowing the formation of a hydrogen bond between Gln130^1.25b^ and Arg23^P^, which is not observed in the LA-PTH–PTH1R–G_s_ complex (18).

Besides the distinct conformations of TMs and ECLs, PTH1R and PTH2R use different amino acids (including Tyr318^5.39b^, Lys197^2.67b^, Arg305^ECL2^ in PTH2R) to recognize their peptides (Fig. 3B). PTH2R uses a polar residue Tyr318^5.39b^ to form hydrogen bonds with Asp7^P^ and Arg11^P^ of TIP39, while PTH1R has a hydrophobic isoleucine (Ile363^5.39b^) at the corresponding site (Fig. 3B). Interestingly, Asp7^P^ is one unique site of TIP39 that corresponding to Ile5^P^ of PTH and His5^P^ of PTHrP (Fig. 3B). In our MD simulations of PTH2R engaging different peptides, Asp7^P^ of TIP39 stably formed hydrogen bonds with Tyr318^5.39b^, while Ile5^P^ of PTH made hydrophobic interactions with Tyr318^5.39b^ (Fig. 3C, Fig. S6). In contrast, no hydrogen bond or hydrophobic interaction between His5^P^ of PTHrP and Tyr318^5.39b^ were observed in the PTHrP-bound PTH2R simulations. This observation is consistent with our mutagenesis studies, where Y318A mutation of PTH2R decreased TIP39 potency by 794-fold but increased PTH potency by 4-fold (Fig. S7). Lys197^2.67b^ of PTH2R has stable hydrophobic interactions with the aromatic Phe10^P^ of TIP39, which is stronger than the interactions with the smaller hydrophobic side chains of corresponding residues at PTH (Met8^P^) or PTHrP (Leu8^P^) (Fig. 3C). TIP39 and PTH share a conserved negatively charged residue (TIP39 Glu21^P^ or PTH Glu19^P^), but PTHrP has a positively charged arginine (Arg19^P^) instead. In the simulations, either Glu21^P^ (TIP39) or Glu19^P^ (PTH) formed putative salt bridges with a positively charged ECL2 residue Arg305^ECL2^ (Fig. 3C), while PTHrP Arg19^P^ repelled Arg305^ECL2^ and might impede the peptide binding. In addition to the residues crucial to ligand specificity, there are several conserved contacts shared by PTH1R and PTH2R. Either Glu4^P^ of LA-PTH or Asp6^P^ of TIP39 contributes salt bridges with Arg^2.60b^ and hydrogen bonds with Tyr^1.47b^. At the middle region of these three peptides, two residues (Ala5^P^/Ala9^P^ in TIP39, Ser3^P^/Leu7^P^ in both PTH and PTHrP) hydrophobically interacted with Leu399^7.43b^ in all simulations (Fig. S6B-D). At the C termini of peptides, a hydrophobic residue (Trp25^P^ in TIP39, Trp23^P^ in PTH and Phe23^P^ in PTHrP) with a large side chain constantly interacts with two ECD residues Ile34^ECD^ and Ile38^ECD^ (Fig. S6H-J). I34A and I38A mutants significantly reduced the potencies of TIP39 and PTH (Fig. S7 and Table S5), which is fully consistent with the simulation results.

**Figure 3.**
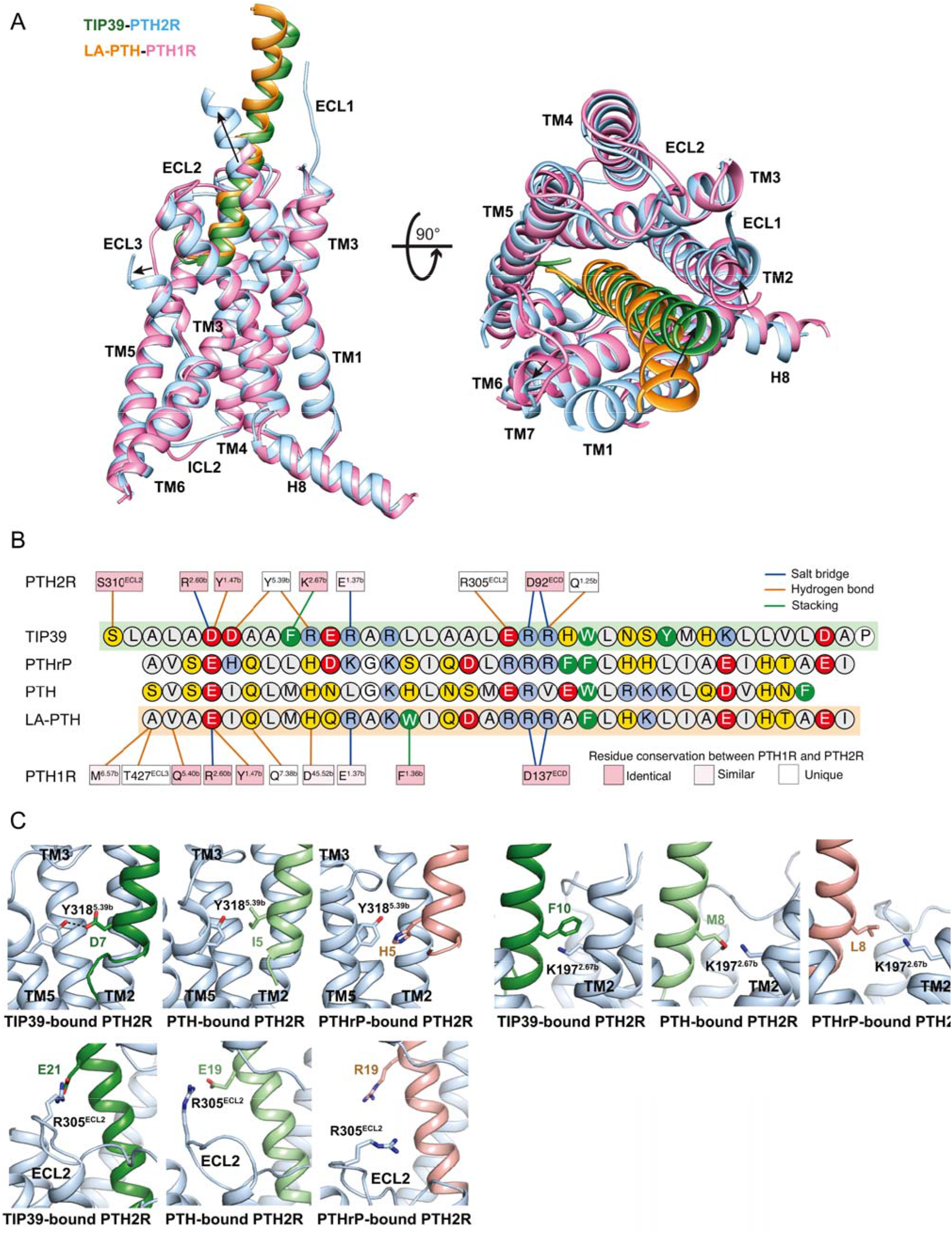
Ligand specificity between PTH1R and PTH2R. **(*A*)** Structural comparison of TIP39-PTH2R-G_s_ and LA-PTH-PTH1R-G_s_ complexes. Receptor ECD and G protein are omitted for clarity. **(*B*)** Schematic diagram of interactions between peptide and receptor. Conserved residues in PTH1R and PTH2R are highlighted in pink, while those similar are shown in light pink. Amino acid residues of peptides are colored: red, negatively charged; blue, positively charged; yellow, hydrophilic; green, aromatic; gray, hydrophobic. Hydrophobic contacts are omitted for clarity. **(*C*)** Representative snapshots from MD simulations showing the key residues that determine the ligand specificity of PTH2R (blue). TIP39, PTH and PTHrP are depicted in green, light green and pink, respectively.

### Antagonism by TIP(7–39)

Deletion of six residues from the N terminus of TIP39 resulted in an antagonist, TIP(7–39) (Fig. 4A) (17). In the MD simulations of TIP(7–39)-bound PTH2R, the receptor spontaneously transitioned from the active conformation to an inactive-like one, displaying a smaller TM6 helix kinking angle (76.9° ± 9.1°) compared with that of TIP39 bound PTH2R (88.2° ± 8.3°) (Fig. 4B-C). In the TIP39-bound PTH2R simulations, the N terminus of the peptide resided between TM5 and TM6 helices (Fig. 4B, D). Particularly, the residues located at the N terminus of TIP39 interacted with TM5 residues (Tyr318^5.39b^, Gln319^5.40b^, Ile322^5.43b^, Leu323^5.44b^ and Ile326^5.47b^), close to the ligand-binding pocket (Fig. 4D, F). Single-point mutations of these residues such as Y318A and Q319A showed significantly reduced cAMP accumulations induced by TIP39 (Fig. 2C), which is consistent with the MD observations. In the TIP(7–39)-bound PTH2R simulations, the interactions between the N terminus of the peptide and TM5 helix were missing (Fig. 4E, F). Consequently, the average backbone distance between TIP(7–39) and TM5 helix was 12.9 ± 0.6 Å, approximately 6 Å longer than that of TIP39-bound PTH2R (7.2 ± 0.7 Å). TIP(7–39) did not have stable interactions with TM6 helix (Fig. 4F). Without direct contacts with the peptide, the C terminus of TM6 helix moved upwards to reduce the kinking (Fig. 4C, E). In the TIP39-bound PTH2R simulations, stable insertion of the N terminus led to a large movement of 9.8 ± 0.8 Å between TM5 and TM6 helices on the extracellular side, which kept the large kinking angle of the TM6 helix (Fig. 4D). In addition, a conserved TM7 residue Gln405^7.49b^ could form hydrogen bonds with the backbone atoms of the TM6 residue Leu370^6.47b^ to further stabilize the kinking conformation of the TM6 helix during the simulations (Fig. 4D, Fig. S8). In the TIP(7–39)-bound PTH2R simulations, however, the polar interactions between TM6 Leu370^6.47b^ and TM7 Gln405^7.49b^ were missing in the receptor core (Fig. 4E, Fig. S8).

**Figure 4.**
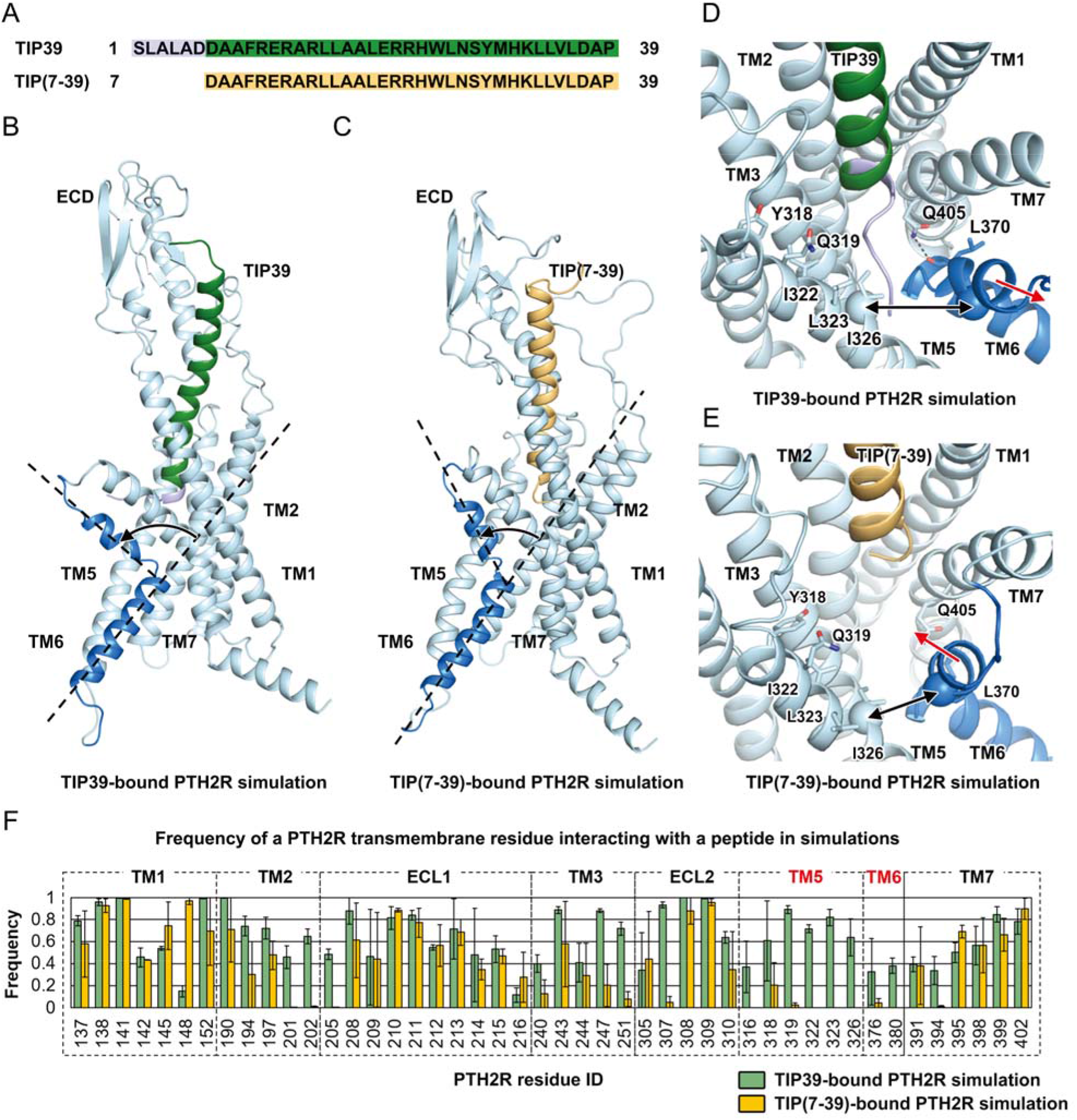
Molecular mechanism of the TIP(7–39) antagonism at PTH2R. **(*A*)** Sequence alignment between TIP39 and TIP(7–39). **(*B*)** A representative snapshot from the TIP39-bound PTH2R simulation system showing a TIP39-induced conformational change of the TM6 helix. **(*C*)** A representative snapshot from the TIP(7–39)-bound PTH2R simulation system showing a TIP(7–39)-induced conformational change of the TM6 helix. **(*D*)** A representative snapshot from the TIP39-bound PTH2R simulation system showing the N terminus of TIP39 insertion between TM5 and TM6 helices. Key residues are shown as sticks. Hydrogen bonds are show as dash lines. The Cα atoms of residues Ile326^5.47b^ and Ile377^6.54b^ are shown as spheres. **(*E*)** A representative snapshot from the TIP(7–39)-bound PTH2R simulation system showing a conformational change of the TM6 helix. **(*F*)** Frequency of a PTH2R residue interacting with TIP39 (green) or TIP(7–39) (yellow) in simulations. The frequency value indicates the stability of a particular residue-peptide interaction. A large interacting frequency indicates a stable interaction.

At the bottom of the ligand-binding pocket, the aspartic acid residue Asp6^P^ of TIP39 was mainly responsible for interacting with Tyr152^1.47b^ and Arg190^2.60b^ (Fig. S8); two residues that govern the functionality of PTH2R (Fig. 2). Multiple hydrogen bonds were formed between these residues. The average atom distances from the Asp6^P^ in TIP39 to Tyr152^1.47b^ and Arg190^2.60b^ were 2.8 ± 0.3 Å and 2.8 ± 0.2 Å, respectively. Because TIP(7–39) does not have Asp6^P^, its Asp7^P^ flipped into the receptor core to interact with Tyr152^1.47b^ and Arg190^2.60b^ instead of Asp6^P^ seen with TIP39 (Fig. S8). Through the C terminus, both of TIP39 and TIP(7–39) are capable of stably interacting with the ECD. Ligand binding patterns at the ECD were almost identical in TIP39 bound and TIP(7–39) bound PTH2R simulations (Fig. S8C). These findings demonstrate that the C terminus of TIP39 and TIP(7–39) contribute to ligand binding, while the N terminus determine receptor activation.

## Disease-associated mutation

Several naturally occurring mutations in PTH2R have been reported to cause multiple hereditary human disorders (10, 11). Of them, two mutations (S158F and G258D) occur in regions that were well-solved in our PTH2R structure, but only G258D significantly affected TIP39 elicited cAMP accumulation (Fig. S9 and Table S6). Gly258^3.51b^ is located at the intracellular side of TM3 helix (a part of the G-protein-binding interface) and implicated in syndromic short stature (10). In the wild-type (WT) PTH2R MD simulations, Gly258^3.51b^ was surrounded by several hydrophobic residues (Leu259^3.52b^, Leu332^5.53b^ and Phe372^6.49b^) of helices TM3, TM5 and TM6 (Fig. 5A, B). Particularly, Gly258^3.51b^ and Phe372^6.49b^ are constantly interacting with an average distance of 3.3 ± 0.2 Å, forming the key helix-helix interface between helices TM3 and TM6. The hydrophobic interactions among Leu259^3.52b^, Leu332^5.53b^ and Phe372^6.49b^ also stabilized the tight bundle of helices TM3, TM5 and TM6 at the G protein-binding interface. The inter-residue distance between any two of these three residues was smaller than 4 Å in the WT simulations. However, in the G258D simulations, Asp258^3.51b^ disrupted the hydrophobic interactions involving Leu259^3.52b^, Leu332^5.53b^ and Phe372^6.49b^ (Fig. 5C, D). The inter-residue distance between any two of the four residues (Asp258^3.51b^, Leu259^3.52b^, Leu332^5.53b^ and Phe372^6.49b^) was larger than 5 Å in the G258D simulations. As a result, the conformations of helices TM3, TM5 and TM6 were interrupted at the intracellular side of the receptor. In the WT simulations, Ile265^3.58b^ and Val339^5.60b^ formed stable hydrophobic interactions to closely pack TM3 and TM5 helices at the intracellular side. However, in the G258D simulations, no direct interactions between these two residues were observed, therefore, the intracellular side of TM3, TM5 and TM6 were distorted and unfavorable to bind to a G-protein (Fig. 5E, F).

**Figure 5.**
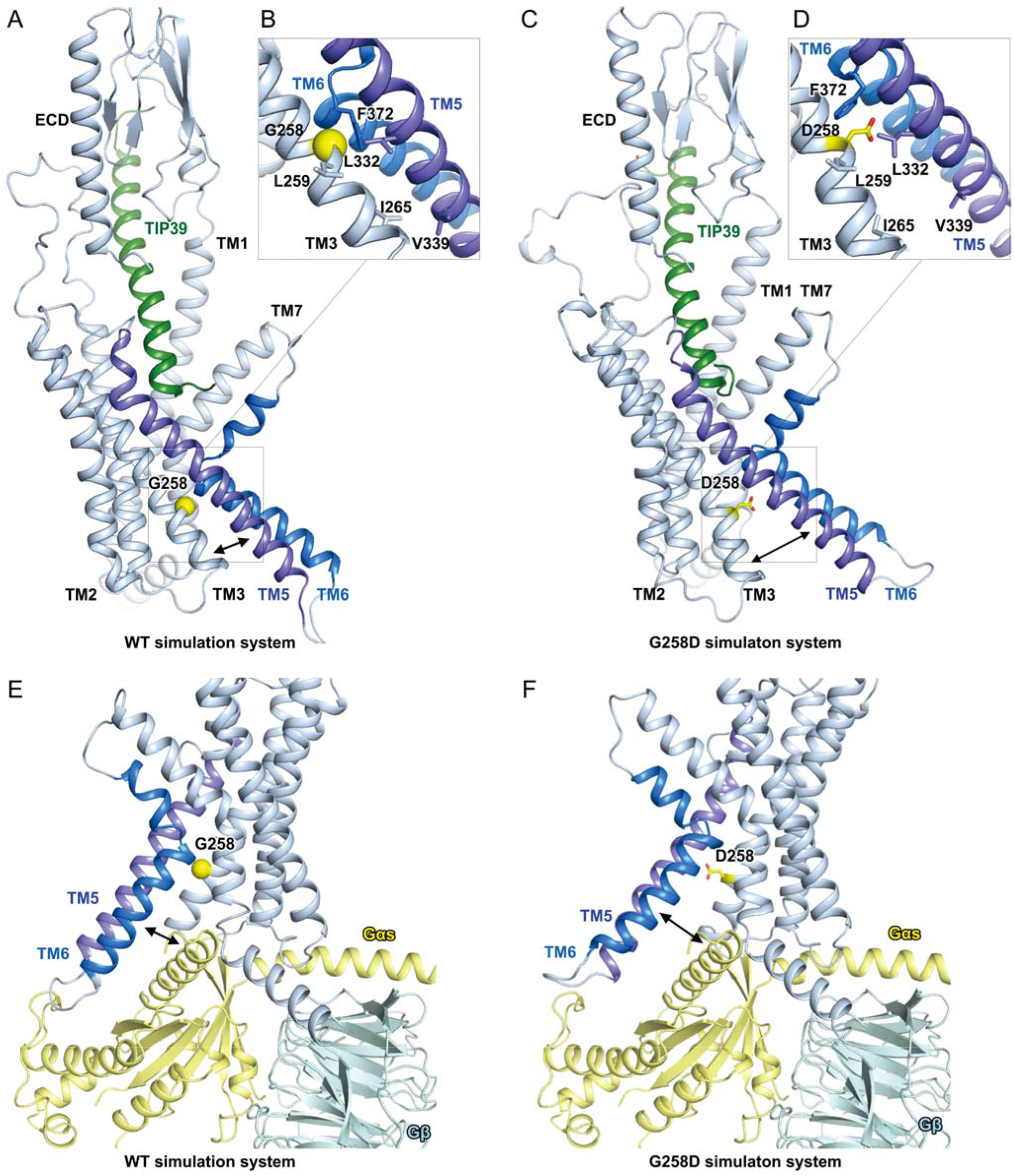
G258D mutation disrupts the G protein-binding interface of PTH2R in MD simulations. **(*A*)** A representative snapshot from the wild-type (WT) PTH2R simulations. **(*B*)** Key interactions stabilizing the helical bundle of TM3, TM5 and TM6 in the WT PTH2R simulations. Key residues are shown as sticks. Gly258^3.51b^ is shown as a yellow sphere. **(*C*)** A representative snapshot from the G258D PTH2R simulations. **(*D*)** Asp258^3.51b^ disrupts the hydrophobic interactions among helices TM3, TM5 and TM6 in the G258D PTH2R simulations. Key residues are shown as sticks. Asp258^3.51b^ is highlighted in yellow. **(*E*)** A representative conformation of the G protein-binding interface of the WT PTH2R in simulations. The cryo-EM structure of TIP39-PTH2R-G_s_ complex was aligned to the simulation resulting conformation to show the position of G protein with respect to the receptor. **(*F*)** A representative conformation of the G protein-binding interface of the G258D PTH2R in simulations.

## Discussion

We used the single-particle cryo-EM approach to solve the high-resolution structure of the TIP39-bound PTH2R in complex with G_s_. It provides essential structural information for understanding how PTH2R recognizes a peptide ligand and couples to G_s_ in the active state. Compared with other class B1 GPCRs (18, 20, 21), PTH2R shows a unique peptide-receptor binding interface, 65% of which is contributed by the N terminus of TIP39. Unlike typical peptides of class B1 GPCRs that adopt helix conformations at their N termini, TIP39 displays a closed loop at the N-terminal (Fig. 2A, Fig. S4). Both cryo-EM and MD simulation data indicate that the unique N terminus of TIP39 not only facilitates a deep insertion of the peptide into the receptor core (Figs. 2–3 and Fig. S6A), but also participates in PTH2R activation via interacting with TM5 and TM6. These findings suggest a possible common mechanism of ligand-induced receptor activation by peptides with loop conformations at the N-terminal (Fig. S6A).

Due to the relatively high-resolution (2.8 Å) of the structure, we were able to address the ligand specificity of PTH2R against three functionally important peptides (TIP39, PTH and PTHrP). Their actions could be divided into three modes: potent (TIP39), mild (PTH) and weak (PTHrP), respectively (4, 13, 16). Integrating MD simulation with mutagenesis studies, we identified key residues responsible for ligand recognition and characterized important receptor-peptide interactions that govern ligand specificity. MD simulations showed that three residues in PTH2R (Lys197^2.67b^, Arg305^ECL2^ and Tyr318^5.39b^) are selective against different peptides. Lys197^2.67b^ stably interacts with Phe10^P^ of TIP39 (Fig. 3C). Arg305^ECL2^ forms putative salt bridges with Glu21^P^ of TIP39 or Glu19^P^ of PTH, but repels Arg19^P^ of PTHrP (Fig. 3C). Tyr318^5.39b^ forms putative hydrogen bonds with Asp7^P^ of TIP39 as well as hydrophobic interactions with Ile5^P^ of PTH, but fails to interact with His5^P^ of PTHrP (Fig. 3C). Notably, Gardella and colleagues have reported that the substitution of Ile5^P^ of PTH with a histidine decreases the peptide potency on PTH2R, but the substitution of His5^P^ of PTHrP with an Isoleucine significantly increases the potency (27), a phenomenon that is highly consistent with our MD simulations.

Activation of class B1 GPCRs is characterized by opening the transmembrane helix bundles at the extracellular side, while the intracellular side undergoes conformational changes to accommodate G protein. Such hourglass-like opening of both extracellular and intracellular portions of the TMD requires TM6 helix to be bended and tightly tethered to the other helices at the receptor core (18, 20, 21). In this work, we link the opposing activities of an agonist (TIP39) and an antagonist TIP(7–39) the TM6 helix of PTH2R: both of them bind to the ECD, with TIP39 inserted into the base of the TMD orthosteric pocket. The N terminus of TIP39 inserts between TM5 and TM6 helices to enhance the kinking of TM6 helix, a step essential to class B1 GPCR activation. A conserved TM7 residue (Gln405^7.49b^) forms hydrogen bonds with the backbone atoms of the TM6 (Leu370^6.47b^) might further stabilize the kinking conformation. A glutamine residue at the corresponding site of other class B1 GPCRs has been reported to act as a molecular switch between receptor activation states (18). Upon full activation of a receptor, this TM7 glutamine reorients downward to establish hydrogen bonds with the TM6 helix residues of a conserved LXXG motif (Fig. S10). This conserved rearrangement of the TM7 residue close to the receptor core enables the stabilization of the distinct kink in TM6 helix to mediate the simultaneous opening of the intracellular and extracellular side. Compared with TIP39, TIP(7–39) has the same C terminus but misses six residues at N terminus. MD simulations revealed that it binds to the ECD via C terminus, but fails to active the receptor due to lack of stable interaction with TM6 helix. These findings disclose the structural basis of PTH2R antagonism and underscore an indispensable role of the N terminus of an agonist in activating PTH2R. This might extend to other class B GPCRs. In fact, N-terminal truncation of PTH, such as PTH(7–34), also results in antagonists for PTH1R (28–30).

Like some other class B1 GPCRs that are implicated in multiple hereditary human disorders (5–10, 12), PTH2R also has several disease-associated mutations, such as the naturally occurring mutation G258D that is associated with syndromic short stature (10). By means of MD simulations, we hypothesize that this mutation might disrupt the active conformation of PTH2R, leading to impaired receptor function (Fig. S9). Surrounded by several hydrophobic residues (Leu259^3.52b^, Leu332^5.53b^ and Phe372^6.49b^) of helices TM3, TM5 and TM6, Gly258^3.51b^ is located nearby the G-protein binding interface of PTH2R. In the simulations of the G258D mutant receptor, Asp258^3.51b^ disturbs the hydrophobic interactions involving Leu259^3.52b^, Leu332^5.53b^ and Phe372^6.49b^ to distort the helical bundle of TM3, TM5 and TM6 at the intracellular side (Fig. 5A-D). Consequentially, the G protein-binding interface is disordered and unfavorable to bind to a heterotrimeric G_s_ protein (Fig. 5E, F). While cAMP response was not affected in S158F mutant (Table S6), both S158F and G258D showed impaired G_q_ coupling (Fig. S11). Based on the atomic-level structural information of PTH2R, we were able to quantitatively interpret the mutational data. This understanding provides valuable information to develop new therapies for disorders associated with PTH2R.

## Materials and Methods

The data that support the findings of this study are available in this paper and/or in supplementary information. Atomic coordinates of the TIP39-PTH2R-G_s_ complex have been deposited in the Protein Data Bank (https://www.rcsb.org/) under accession code 7F16. The electron microscopy maps have been deposited in the Electron Microscopy Data Bank (EMDB) under accession number EMD-31405.

### Construct

The human PTH2R (residues 25-442) was cloned into the pFastBac vector (Invitrogen) with its native signal peptide replaced by haemagglutinin (HA) signal peptide to enhance receptor expression. LgBiT subunit (Promega) was fused at the C-terminus of PTH2R connected by a 20-amino acid linker. A TEV protease cleavage site and double maltose-binding protein (2MBP) tag were fused after LgBiT subunit. A dominant-negative human Gαs (S54N, G226A, E268A, N271K, K274D, R280K, T284D and I285T) (31) was generated to stabilize the interaction with the βγ subunits. A 15-amino acid linker and SmBiT subunit (peptide 86, Promega) were attached to the C-terminus of rat Gβ1. Human DNGαs, rat Gβ1 and bovine Gγ2 were cloned into pFastBac vector, respectively.

### TIP39-PTH2R-G_s_ complex formation and purification

After dounce homogenization of High Five insect cell pellets in lysis buffer (20 mM HEPES, pH 7.4, 100 mM NaCl, 10% (v/v) glycerol supplemented with EDTA-free protease inhibitor cocktail, Topscience), membrane was collected at 65,000× *g* for 35 min and homogenized again in lysis buffer. The complex formation was initiated by addition of 20 μM TIP39 (GL Biochem), 15 μg/mL Nb35, 25 mU/mL apyrase (NEB), 5 mM CaCl_2_, 10 mM MgCl_2_, 1 mM MnCl_2_ and 100 μM TCEP for 1.5 h incubation at room temperature (RT). The membrane was solubilized by 0.5% (w/v) lauryl maltose neopentyl glycol (LMNG; Anatrace) and 0.1% (w/v) cholesterol hemisuccinate (CHS; Anatrace) for 2 h at 4°C. After centrifugation at 65,000× *g* for 35 min, the supernatant was separated and incubated with amylose resin (NEB) for 2 h at 4°C. The resin was collected and packed into a gravity flow column and washed with 5 column volumes of 5 μM TIP39, 0.1% (w/v) LMNG, 0.02% (w/v) CHS, 20 mM HEPES, pH7.4, 100 mM NaCl, 10% (v/v) glycerol, 5 mM MgCl_2_, 1 mM MnCl_2_ and 25 μM TCEP, followed by 20 column volumes of washing buffer with decreased concentrations of detergents 0.03% (w/v) LMNG, 0.01% (w/v) GDN and 0.008% (w/v) CHS. 2MBP-tag was removed by His-tagged TEV protease (home-made) during overnight incubation. The complex was concentrated using an Amicon Ultra Centrifugal filter (MWCO, 100 kDa) and subjected to a Superose 6 Increase 10/300 GL column (GE Healthcare) that was pre-equilibrated with running buffer containing 20 mM HEPES, pH 7.4, 100 mM NaCl, 2 mM MgCl_2_, 100 μM TCEP, 5 μM TIP39, 0.00075% (w/v) LMNG, 0.00025% (w/v) GDN and 0.0002% (w/v) CHS. Eluted fractions containing the TIP39-PTH2R-G_s_ complex were pooled and concentrated. All procedures mentioned above were performed at 4°C.

### Cryo-EM data acquisition

The purified TIP39–PTH2R–Gs complex (3 μL at 8.5 mg per mL) was applied on a glow-discharged holey carbon grid (Quantifoil R1.2/1.3). Vitrification was performed using a Vitrobot Mark IV (ThermoFisher Scientific) at 100% humidity and 4°C. Cryo-EM imaging was processed on a Titan Krios (FEI) equipped with a Gatan K3 Summit direct electron detector in the Center of Cryo-Electron Microscopy, Shanghai Institute of Materia Medica, CAS (China). The microscope was operated at 300 kV accelerating voltage, at a nominal magnification of 95,694× in counting mode, corresponding to a pixel size of 0.5225 Å. In total, 3614 movies were obtained.

### Model building and refinement

Cryo-EM structure model of the PTH2R–G_s_–Nb35 complex was built using the cryo-EM structure of PTH1R–G_s_–Nb35 (PDB code: 6NBF) as initial model. The model was docked into the EM density map using Chimera (32), followed by iterative manual adjustment and rebuilding in COOT (33). Real space refinement was performed using Phenix (34). The model statistics were validated using MolProbity (35). Structural figures were prepared in Chimera and PyMOL (https://pymol.org/2/). The final refinement statistics are provided in Table S1.

### cAMP accumulation assay

The wild-type or mutant PTH2Rs were cloned into pcDNA3.1 vector (Invitrogen) for functional studies. cAMP signal was detected by LANCE cAMP kit (PerkinElmer) according to manufacturer’s instructions. Briefly, HEK-293T cells were seeded onto 6-well culture plates and transiently transfected with different PTH2R constructs using Lipofectamine 2000 transfection reagent (Invitrogen). After 24 h, cells were digested with 0.02% (w/v) EDTA and resuspended by HBSS supplemented with 5 mM HEPES, 0.5 mM IBMX and 0.1% (w/v) BSA, pH 7.4 before seeding onto 384-well microtiter plates (3,000 cells per well). Increased concentrations of TIP39 or PTH (1–34) (1 pM - 1 µM) were used to stimulate transfected cells for 40 min at RT. Eu-tracer and ULight-anti-cAMP working solutions were added to the microtiter plates following 1 h incubation at RT. Fluorescence signals were measured at 620 nm and 650 nm by an EnVision multilabel plate reader (PerkinElmer).

### FITC-labelled ligand binding assay

Competitive binding of TIP39-FITC (GL Biochem) to PTH2R was assessed as described previously (36). Briefly, 24 h after transfection with PTH2R (25-550) or PTH2R (25-442)-20AA-LgBiT, HEK-293□T cells were harvested using 0.2% (w/v) EDTA. They (1 × 10^6^ cells/mL) then mixed with 0.2 μM TIP39-FITC on ice in the dark for 1 h. Seven decreasing concentrations of unlabeled peptide were added and competitively reacted with the cells in binding buffer (HBSS supplemented with 0.5% (w/v) BSA and 20 mM HEPES, pH 7.4) on ice for 2 h. For each sample, 30,000 cells were analyzed for mean fluorescence intensity (with excitation and emission wavelengths of 488 and 518 nm) on a FACScan flow cytometer (ACEA Biosciences), with debris excluded by forward versus side scatter (FSC *vs*. SSC) gating.

### Molecular dynamics simulation

All peptide-bound PTH2R complex models were built based on the TIP39-PTH2R-G_s_ complex structure using Modeller (37). The default parameters were employed to construct the models. The missing backbone and side chains were added. The models with the lowest root mean square deviations from their template structures were selected. To build a simulation system, we placed the complex model into a 1-palmitoyl-2-oleoyl-sn-glycero-3-phosphocholine lipid bilayer. The lipid embedded complex model was solvated in a periodic boundary condition box (95 Å × 95 Å × 170 Å) filed with TI3P water molecules and 0.15 M KCl using CHARMM-GUI (38). Each system was replicated to perform two independent simulations. On the basis of the CHARMM36m all-atom force field (39–41), molecular dynamics simulations were conducted using GROMACS 5.1.4 (42, 43). Further details are provided in supplementary information.

## Acknowledgments

We thank Chenyao Li and Wen Sun for technical assistance. This work was partially supported by National Natural Science Foundation of China 81872915 and 82073904 (M.-W.W.), 32071203 (L.H.Z), 81773792 (D.Y.), 81973373 (D.Y.) and 21704064 (Q.Z.); National Science and Technology Major Project of China – Key New Drug Creation and Manufacturing Program 2018ZX09735–001 (M.-W.W.), 2018ZX09711002–002–005 (D.Y.) and 2018ZX09711002–002–003 (X.C.); the National Key Basic Research Program of China 2018YFA0507000 (M.-W.W.); Ministry of Science and Technology of China 2018YFA0507002 (H.E.X.); Shanghai Municipal Science and Technology Major Project 2019SHZDZX02 (H.E.X.); the Strategic Priority Research Program of Chinese Academy of Sciences XDB37030103 (H.E.X.); Novo Nordisk-CAS Research Fund grant NNCAS-2017–1-CC (D.Y.); Shanghai Science and Technology Development Fund 18ZR1447800 (L.H.Z.), The Young Innovator Association of CAS 2018325 (L.H.Z.) and SA-SIBS Scholarship Program (L.H.Z. and D.Y.). The Youth Innovation Promotion Association of CAS 2018319 (X.C). The cryo-EM data were collected at Cryo-Electron Microscopy Research Center, Shanghai Institute of Materia Medica.

## Author contributions

X.W., L.H.Z. and C.Y.Y. designed the expression constructs, purified the receptor complexes, prepared cryo-EM grids and collected data towards the structure; Y.Z.W. and J.C. developed ligand binding assay; X.Y.Z. and T.X. made map calculation; X.C., L.H.Z. and Q.T.Z. performed structural analysis and prepared figures; X.C. and H.L.J. conducted MD simulations; A.T.D., Y.Z.W. and X.W. conducted functional experiments; D.Y. supervised mutagenesis and signaling studies; H.E.X. and M.-W.W. initiated the project and supervised the project. X.W., X.C., L.H.Z. and M.-W.W. wrote the manuscript with inputs from all co-authors.

## References

1. K. Pal, K. Melcher, H. E. Xu, Structure and mechanism for recognition of peptide hormones by Class B G-protein-coupled receptors. Acta Pharmacol Sin 33, 300–311 (2012).

2. D. Wootten, L. J. Miller, Structural basis for allosteric modulation of class b g protein-coupled receptors. Annu Rev Pharmacol Toxicol 60, 89–107 (2020).

3. H. Juppner et al., A G protein-linked receptor for parathyroid hormone and parathyroid hormone-related peptide. Science 254, 1024–1026 (1991).

4. T. B. Usdin, C. Gruber, T. I. Bonner, Identification and functional expression of a receptor selectively recognizing parathyroid hormone, the PTH2 receptor. J Biol Chem 270, 15455–15458 (1995).

5. E. L. Dimitrov, J. Kuo, K. Kohno, T. B. Usdin, Neuropathic and inflammatory pain are modulated by tuberoinfundibular peptide of 39 residues. Proc Natl Acad Sci U S A 110, 13156–13161 (2013).

6. A. Dobolyi, H. Ueda, H. Uchida, M. Palkovits, T. B. Usdin, Anatomical and physiological evidence for involvement of tuberoinfundibular peptide of 39 residues in nociception. Proc Natl Acad Sci U S A 99, 1651–1656 (2002).

7. E. Sato et al., Activation of parathyroid hormone 2 receptor induces decorin expression and promotes wound repair. J Invest Dermatol 137, 1774–1783 (2017).

8. T. Varga et al., Paralemniscal TIP39 is induced in rat dams and may participate in maternal functions. Brain Struct Funct 217, 323–335 (2012).

9. E. Sato et al., The parathyroid hormone second receptor pth2r and its ligand tuberoinfundibular peptide of 39 residues tip39 regulate intracellular calcium and influence keratinocyte differentiation. J Invest Dermatol 136, 1449–1459 (2016).

10. I. Meulenbelt et al., Strong linkage on 2q33.3 to familial early-onset generalized osteoarthritis and a consideration of two positional candidate genes. Eur J Hum Genet 14, 1280–1287 (2006).

11. D. Tiosano et al., Mutations in PIK3C2A cause syndromic short stature, skeletal abnormalities, and cataracts associated with ciliary dysfunction. PLoS Genet 15, e1008088 (2019).

12. L. Cuttler, Safety and efficacy of growth hormone treatment for idiopathic short stature. J Clin Endocrinol Metab 90, 5502–5504 (2005).

13. T. B. Usdin, S. R. Hoare, T. Wang, E. Mezey, J. A. Kowalak, TIP39: a new neuropeptide and PTH2-receptor agonist from hypothalamus. Nat Neurosci 2, 941–943 (1999).

14. T. J. Gardella, J. P. Vilardaga, International union of basic and clinical pharmacology. xciii. the parathyroid hormone receptors--family b g protein-coupled receptors. Pharmacol Rev 67, 310–337 (2015).

15. B. Z. Leder et al., Effects of abaloparatide, a human parathyroid hormone-related peptide analog, on bone mineral density in postmenopausal women with osteoporosis. J Clin Endocrinol Metab 100, 697–706 (2015).

16. T. B. Usdin, A. Dobolyi, H. Ueda, M. Palkovits, Emerging functions for tuberoinfundibular peptide of 39 residues. Trends Endocrinol Metab 14, 14–19 (2003).

17. S. R. J. Hoare, T. B. Usdin, Tuberoinfundibular peptide (7-39) [TIP(7-39)], a novel, selective, high-affinity antagonist for the parathyroid hormone-1 receptor with no detectable agonist activity. J Pharmacol Exp Ther 295, 761–770 (2000).

18. L. H. Zhao et al., Structure and dynamics of the active human parathyroid hormone receptor-1. Science 364, 148–153 (2019).

19. D. Wootten, J. Simms, L. J. Miller, A. Christopoulos, P. M. Sexton, Polar transmembrane interactions drive formation of ligand-specific and signal pathway-biased family B G protein-coupled receptor conformations. Proc Natl Acad Sci U S A 110, 5211–5216 (2013).

20. X. Zhang et al., Differential GLP-1R binding and activation by peptide and non-peptide agonists. Mol Cell 80, 485–500 e487 (2020).

21. A. Qiao et al., Structural basis of Gs and Gi recognition by the human glucagon receptor. Science 367, 1346–1352 (2020).

22. L. H. Zhao et al., Differential requirement of the extracellular domain in activation of class B G protein-coupled receptors. J Biol Chem 291, 15119–15130 (2016).

23. Y. L. Liang et al., Phase-plate cryo-EM structure of a class B GPCR-G-protein complex. Nature 546, 118–123 (2017).

24. Y. Zhang et al., Cryo-EM structure of the activated GLP-1 receptor in complex with a G protein. Nature 546, 248–253 (2017).

25. Y. L. Liang et al., Phase-plate cryo-EM structure of a biased agonist-bound human GLP-1 receptor-Gs complex. Nature 555, 121-+ (2018).

26. S. Ma et al., Molecular basis for hormone recognition and activation of corticotropin-releasing factor receptors. Mol Cell 77, 669–680 e664 (2020).

27. T. J. Gardella, G. S. Jensen, M. Luck, T. B. Usdin, H. Juppner, Converting parathyroid hormone-related peptide (PTHrP) into a potent PTH-2 receptor agonist. J Bone Miner Res 11, 77–77 (1996).

28. R. W. Cheloha, T. Watanabe, T. Dean, S. H. Gellman, T. J. Gardella, Backbone Modification of a parathyroid hormone receptor-1 antagonist/inverse agonist. Acs Chemical Biology 11, 2752–2762 (2016).

29. S. H. Doppelt et al., Inhibition of the invivo parathyroid hormone-mediated calcemic response in rats by a synthetic hormone antagonist. P Natl Acad Sci USA 83, 7557–7560 (1986).

30. N. Horiuchi, M. F. Holick, J. T. Potts, M. Rosenblatt, A Parathyroid-hormone inhibitor invivo - design and biological evaluation of a hormone analog. Science 220, 1053–1055 (1983).

31. Y. L. Liang et al., Dominant negative G proteins enhance formation and purification of agonist-GPCR-G protein complexes for structure determination. ACS Pharmacol Transl Sci 1, 12–20 (2018).

32. E. F. Pettersen et al., UCSF Chimera--a visualization system for exploratory research and analysis. J Comput Chem 25, 1605–1612 (2004).

33. P. Emsley, K. Cowtan, Coot: model-building tools for molecular graphics. Acta Crystallogr D Biol Crystallogr 60, 2126–2132 (2004).

34. P. D. Adams et al., PHENIX: a comprehensive Python-based system for macromolecular structure solution. Acta Crystallogr D Biol Crystallogr 66, 213–221 (2010).

35. V. B. Chen et al., MolProbity: all-atom structure validation for macromolecular crystallography. Acta Crystallogr D Biol Crystallogr 66, 12–21 (2010).

36. H. Fan et al., The non-peptide GLP-1 receptor agonist WB4-24 blocks inflammatory nociception by stimulating beta-endorphin release from spinal microglia. Br J Pharmacol 172, 64–79 (2015).

37. A. Sali, T. L. Blundell, Comparative protein modelling by satisfaction of spatial restraints. J Mol Biol 234, 779–815 (1993).

38. E. L. Wu et al., CHARMM-GUI membrane builder toward realistic biological membrane simulations. J Comput Chem 35, 1997–2004 (2014).

39. O. Guvench et al., CHARMM additive all-atom force field for carbohydrate derivatives and its utility in polysaccharide and carbohydrate-protein modeling. J Chem Theory Comput 7, 3162–3180 (2011).

40. J. Huang et al., CHARMM36m: an improved force field for folded and intrinsically disordered proteins. Nat Methods 14, 71–73 (2017).

41. A. D. MacKerell et al., All-atom empirical potential for molecular modeling and dynamics studies of proteins. J Phys Chem B 102, 3586–3616 (1998).

42. B. Hess, C. Kutzner, D. van der Spoel, E. Lindahl, GROMACS 4: algorithms for highly efficient, load-balanced, and scalable molecular simulation. J Chem Theory Comput 4, 435–447 (2008).

43. D. Van Der Spoel et al., GROMACS: fast, flexible, and free. J Comput Chem 26, 1701–1718 (2005).

