## Supplemental information for "Molecular insights into differentiated ligand recognition of the human parathyroid hormone receptor 2"

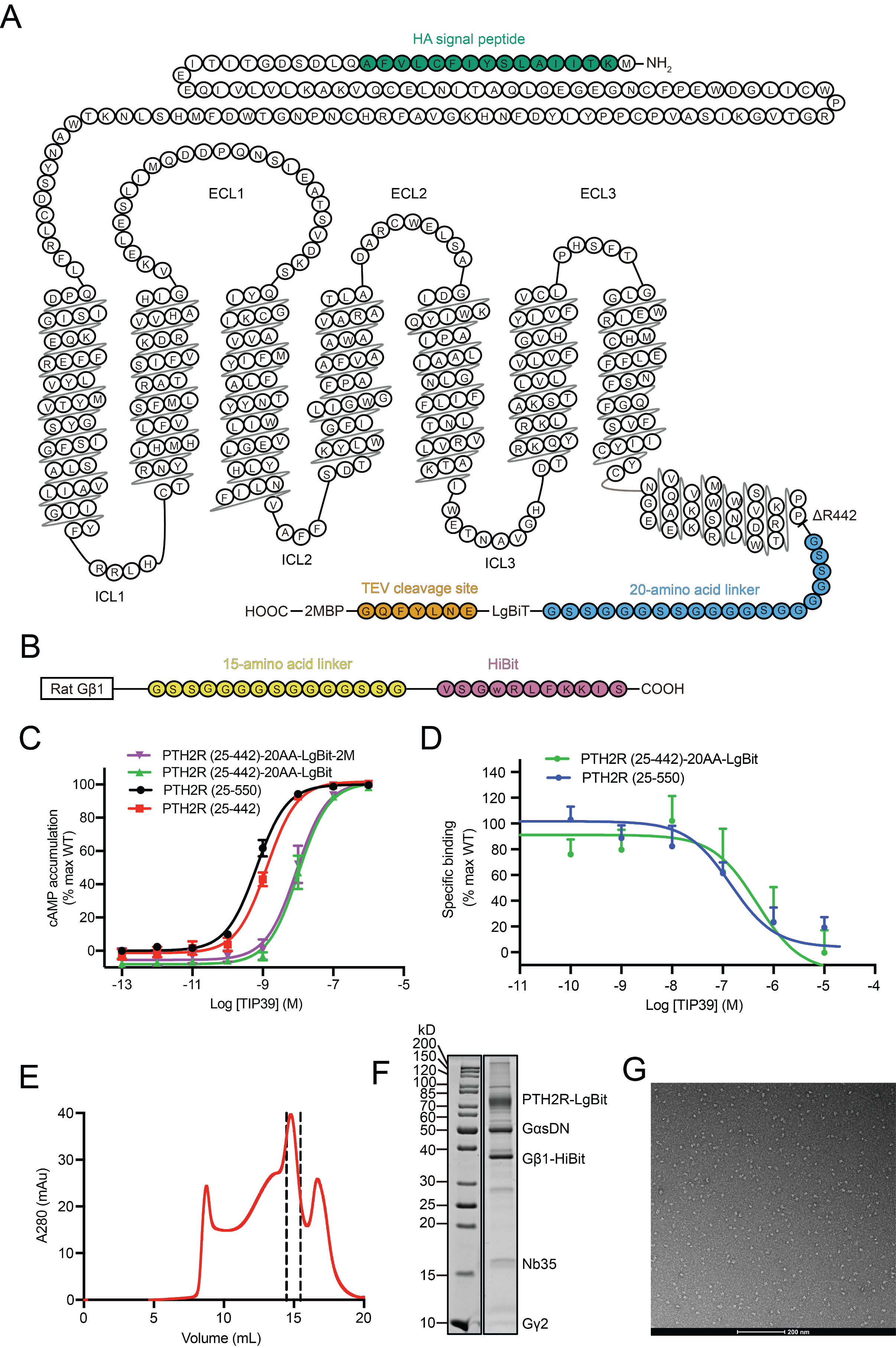

**Fig S1. Purification and characterization of the TIP39–PTH2R–G_s_ complex. (*A*)** Snake representation of human PTH2R construct used in this study. **(*B*)** Rat Gβ1 was attached to HiBiT with a 15-amino acid (15AA) linker between them. **(*C*)** cAMP accumulation in wild-type (WT) and truncated PTH2R expressing cells following cognate ligand stimulation. **(*D*)** FITC-labelled ligand binding properties of WT and truncated PTH2 receptors. **(*E*-*G*)** Analytical size-exclusion chromatography, SDS-PAGE/Coomassie blue stain and representative negative staining image of the purified TIP39-PTH2R-G_s_ complex.

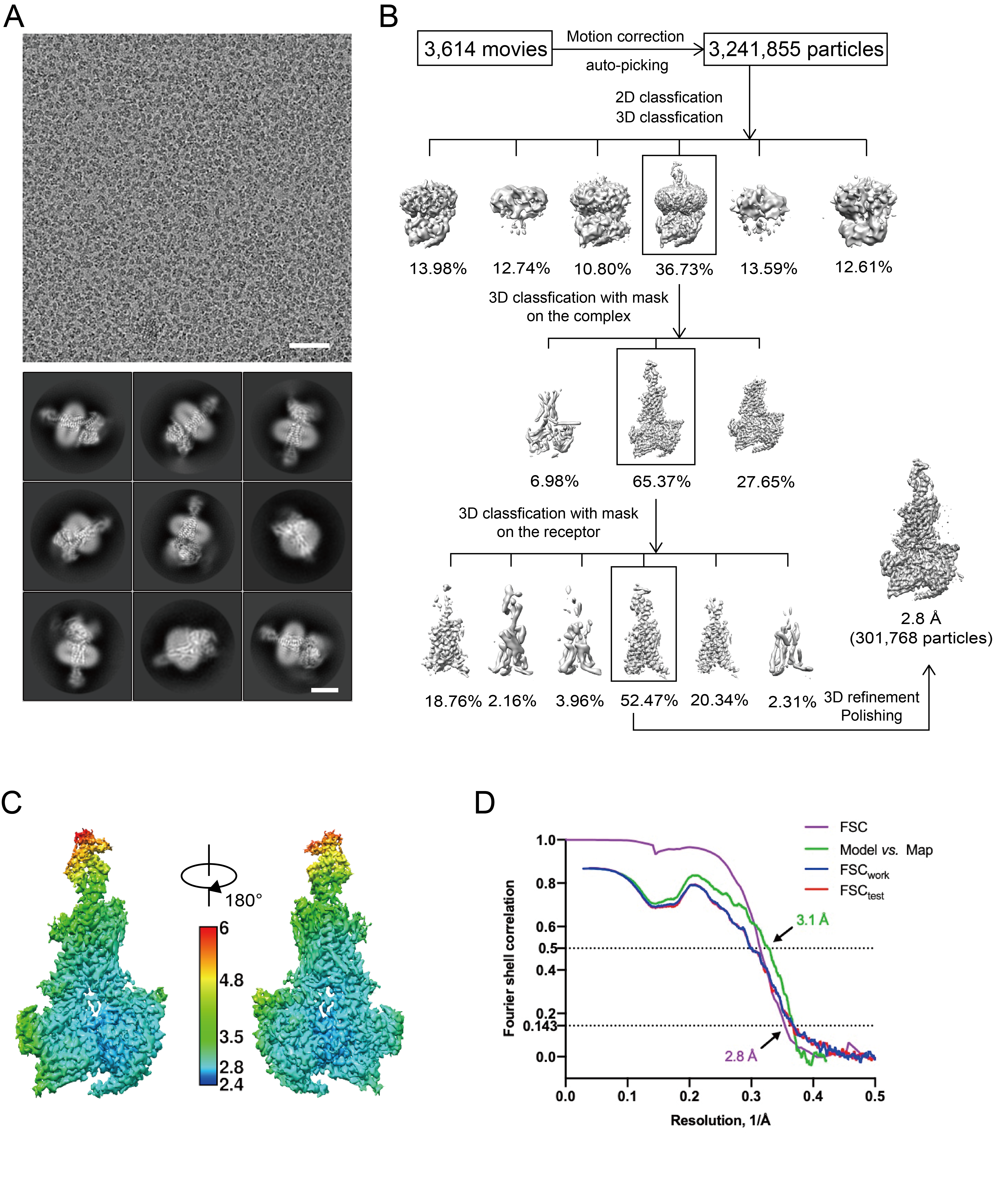

**Fig. S2. Cryo-EM analysis of the TIP39-PTH2R-G_s_ complex. (*A*)** Representative cryo-EM micrographs of the TIP39-PTH2R-G_s_ complex (scale bar: 30 nm). Representative 2D class averages showing distinct secondary structure features from different views (scale bar: 5 nm). **(*B*)** Workflow of cryo-EM data processing for the TIP39-PTH2R-G_s_ complex. **(*C*)** Cryo-EM maps are colored by local resolution (Å). **(*D*)** ‘Gold-standard’ FSC curves of the TIP39-PTH2R-G_s_ complex, indicating the resolution is 2.8 Å at the FSC=0.143, respectively.

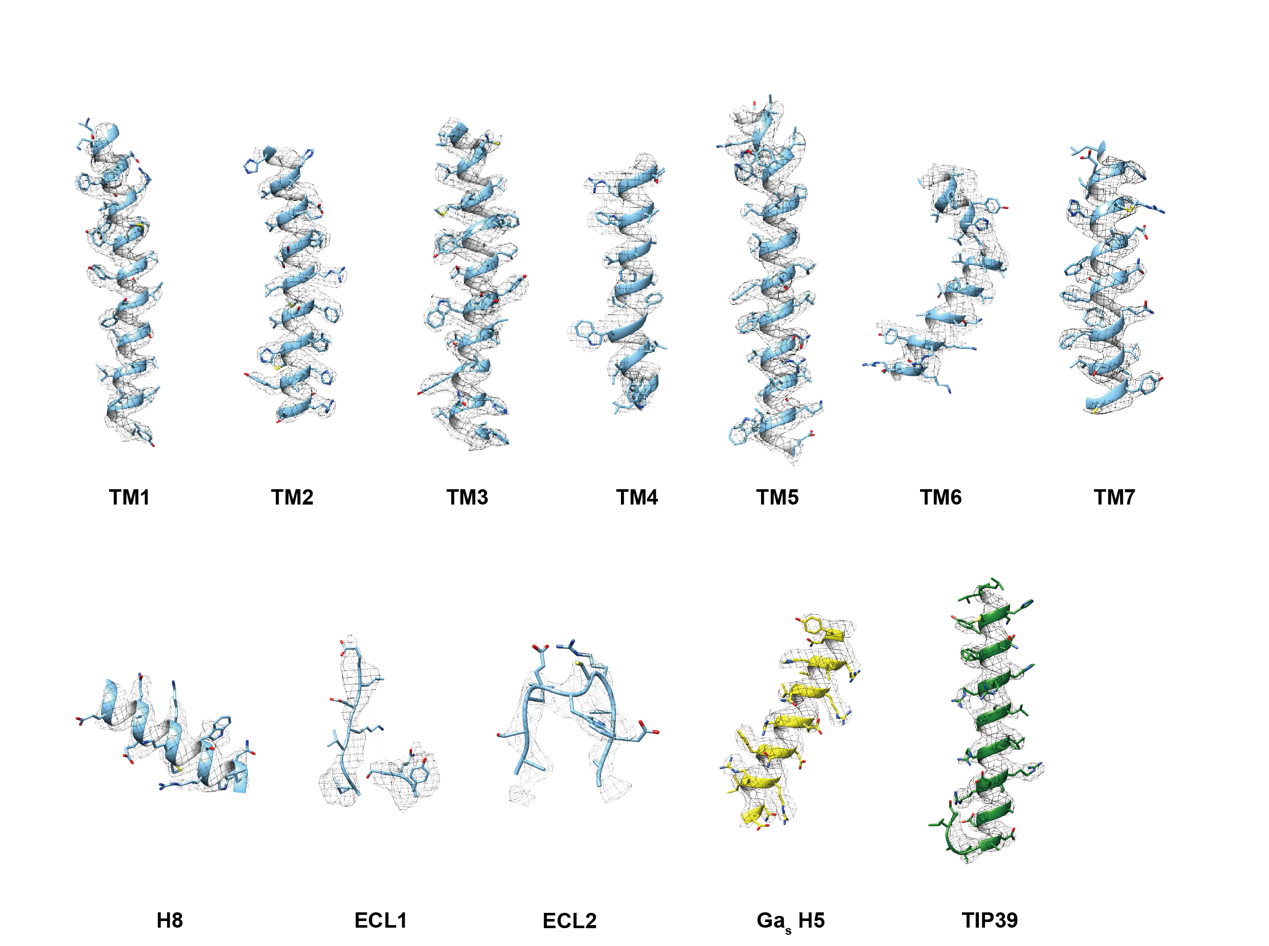

**Fig. S3. Atomic resolution model of the TIP39-PTH2R-G_s_ complex.** EM density map and model are shown for all seven transmembrane α helices, helix 8, ECL1 and ECL2 of PTH2R, the α5 helix of the Gα_s_ Ras-like domain and TIP39.

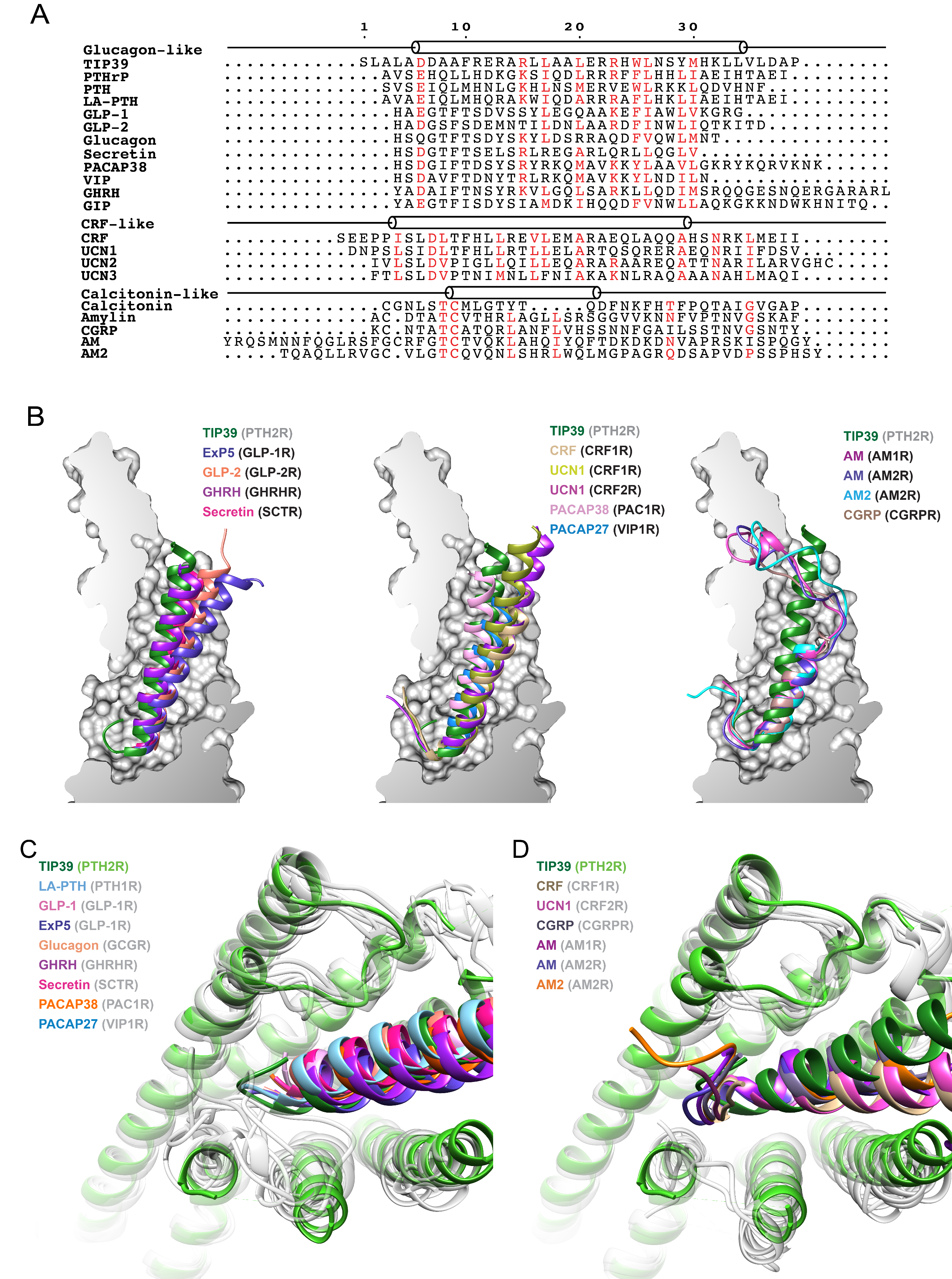

**Fig. S4. Diverse conformations of peptides in the ligand-binding pocket of PTH2R. (*A*)** Structure-based sequence alignment of class B1 GPCR peptide hormones. Residue numbers are shown based on TIP39. The helical structures of peptides are shown as cylinders. **(*B*)** The peptides of class B1 GPCRs in the ligand-binding pocket. Peptides are compared with TIP39 within the PTH2R ligand-binding pocket. **(*C*)** Class B1 GPCR peptide agonists without N terminus loop. **(*D*)** Class B1 GPCR peptide agonists with N terminus loop. Receptor ECD and G protein are omitted for clarity.

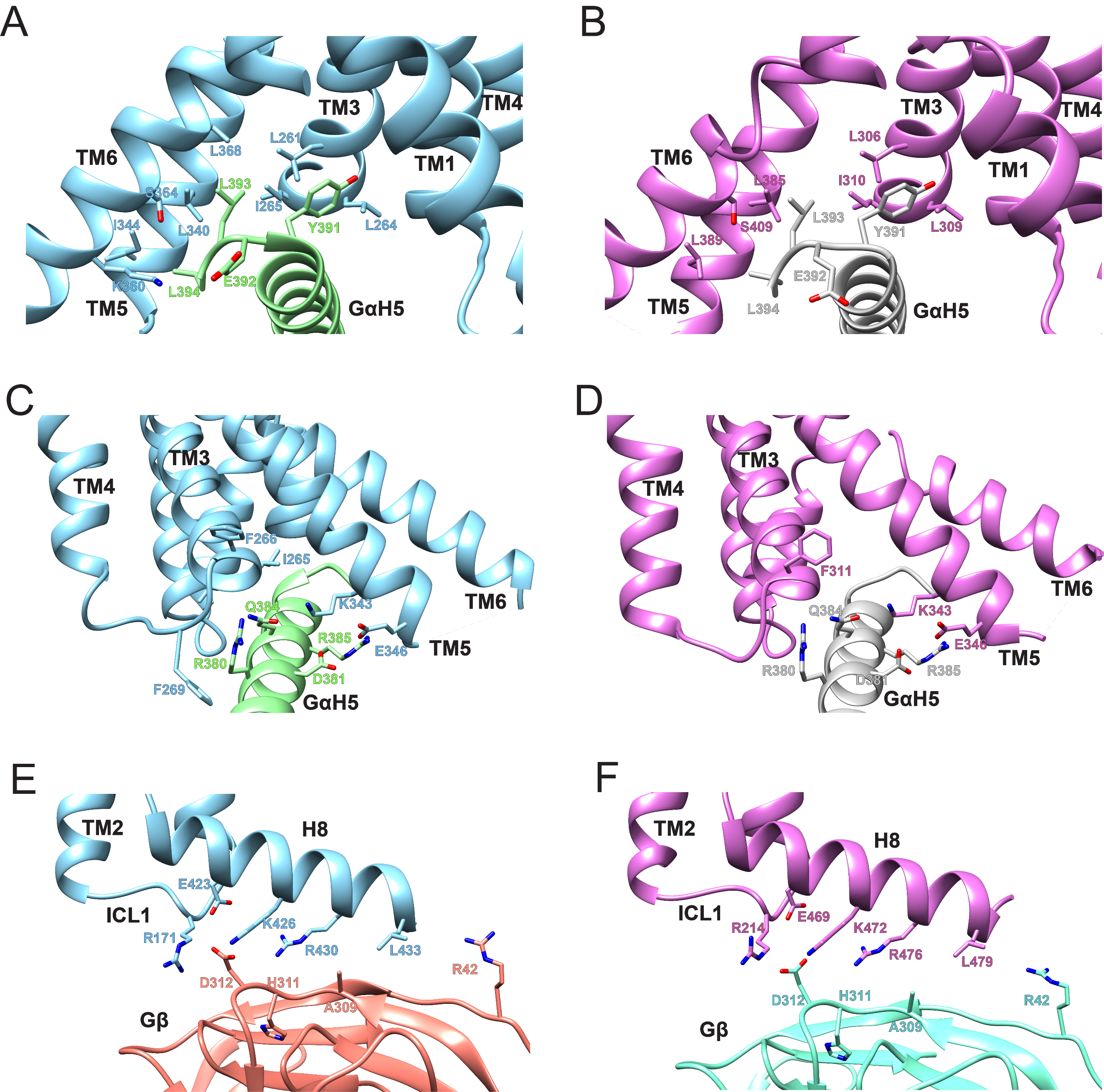

**Fig. S5. G protein-binding interface of PTH2R. (*A*, *C*)** The α5 helix (GαH5) of the Gα_s_ Ras-like domain inserts into an intracellular crevice of PTH2R TMD. **(*E*)** The interface of helix 8 (H8) of PTH2R and Gβ. PTH2R in blue, GαH5 in green, Gβ in salmon. **(*B*, *D*)** The α5 helix (GαH5) of the Gα_s_ Ras-like domain inserts into an intracellular crevice of PTH1R TMD. **(*F*)** The interface of helix 8 (H8) of PTH1R and Gβ. PTH1R in orchid, GαH5 in gray, Gβ in lightcyan.

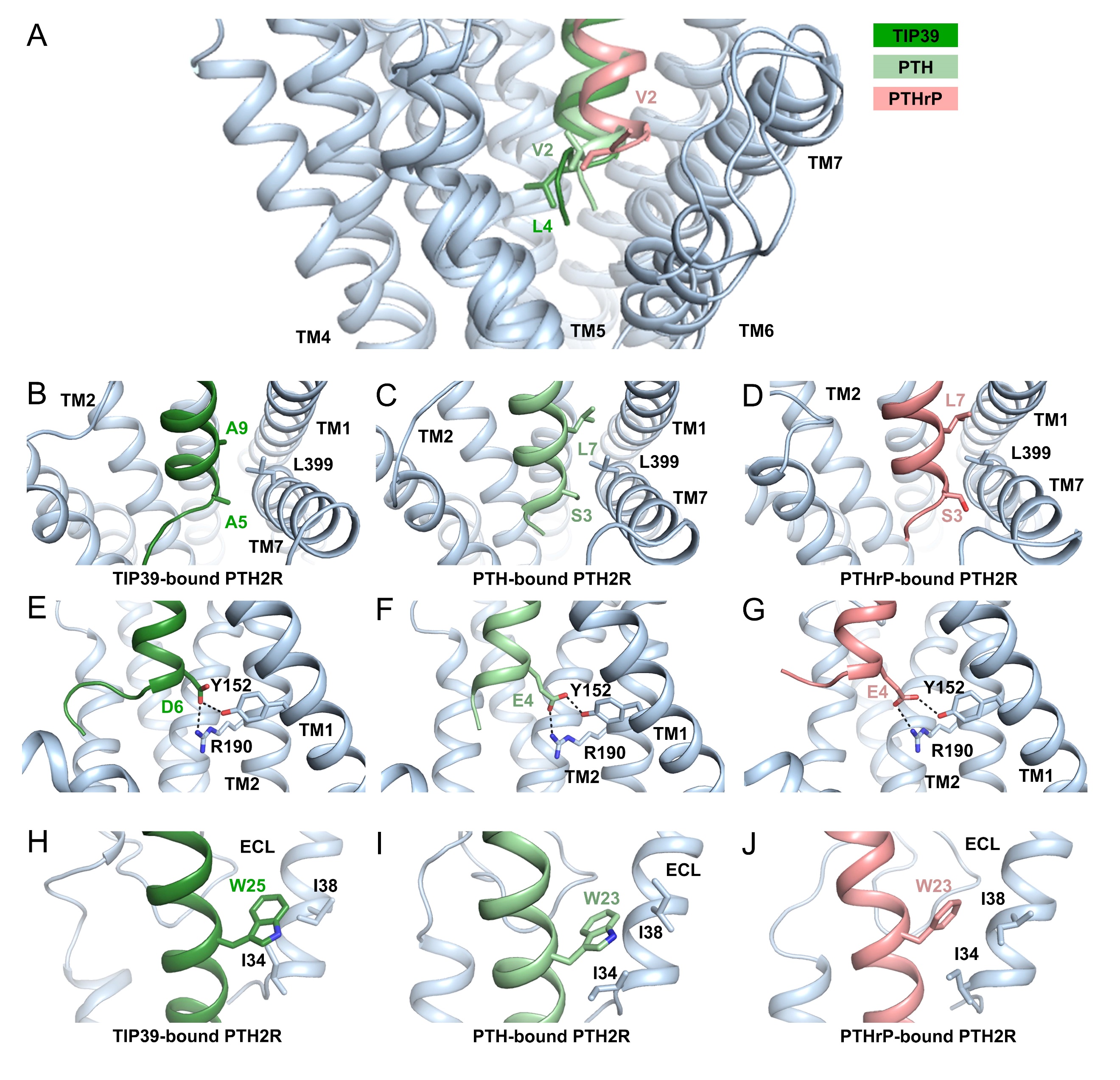

**Fig. S6. Key interactions in the ligand-binding pocket of PTH2R. (*A*)** Representative snapshots from MD simulations showing different peptides inserting into the ligand-binding pocket of PTH2R (*blue*). TIP39 is depicted in green cartoon, PTH is in light green, PTHrP is in pink. Key residues are shown as sticks. **(*B-J*)** Representative snapshots from MD simulations showing key residues responsible for ligand recognition by PTH2R (*blue*). TIP39 is depicted in green cartoon, PTH is in light green, PTHrP is in pink. Key residues are shown as sticks. Salt bridges and hydrogen bonds are showed as dash lines.

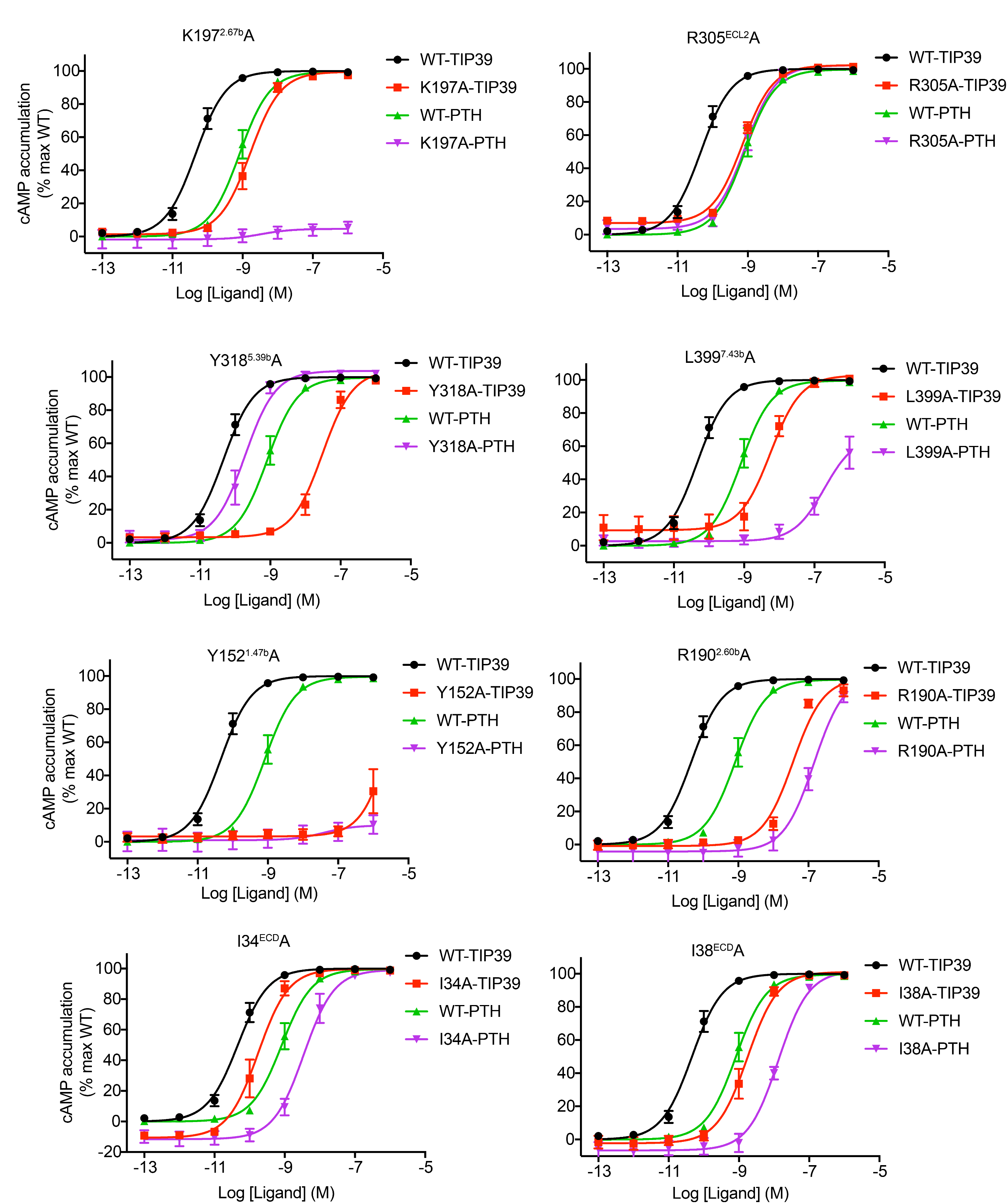

**Fig. S7. Signaling profiles of TIP39 and PTH on the PTH2R mutants.** cAMP accumulation in wild-type (WT) and (single-point) mutant PTH2R expressing cells. Concentration-response curves were generated and graphed as means ± S.E.M. from three independent experiments.

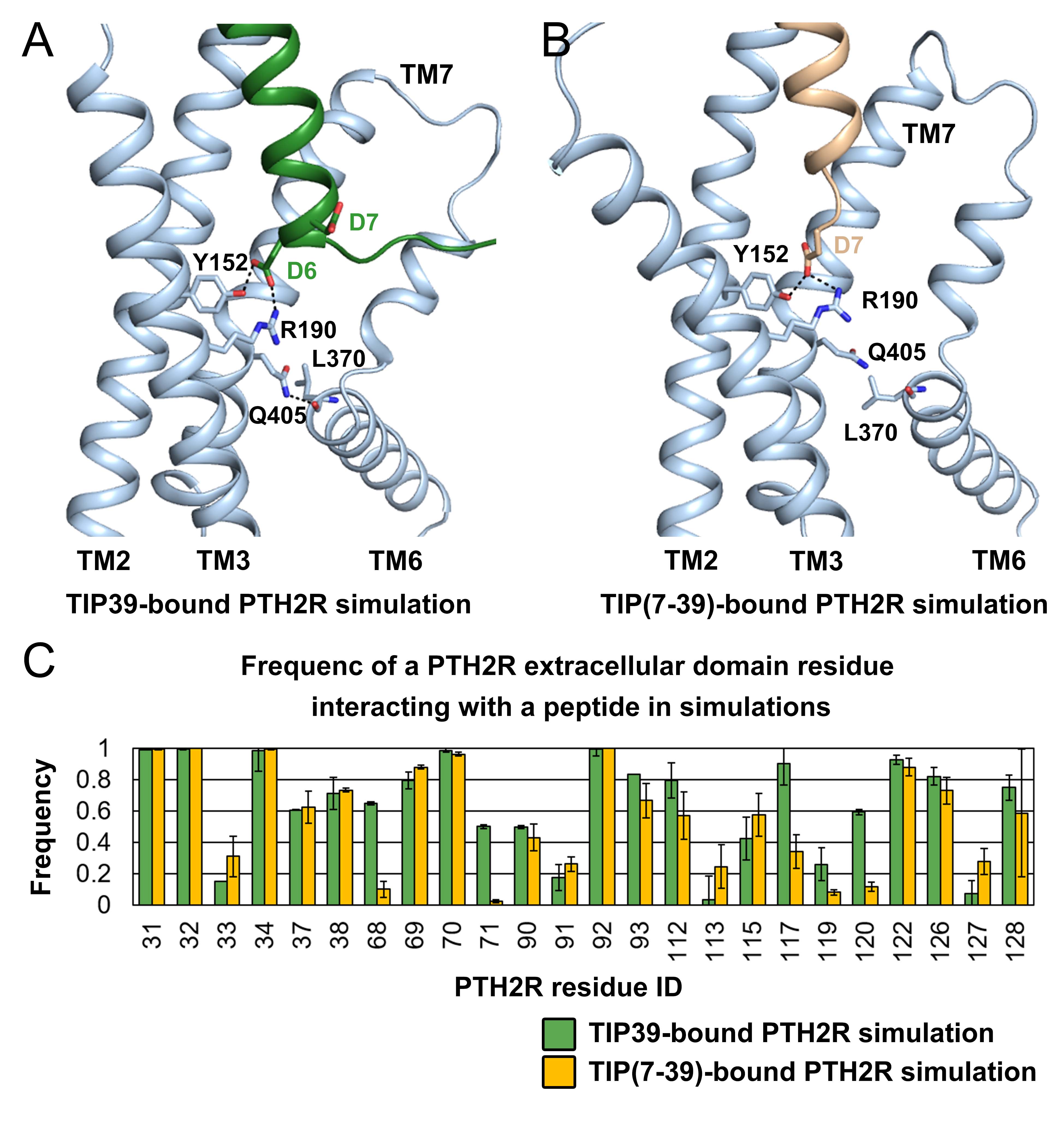

**Fig. S8. Key residues of PTH2R binding to TIP39 and TIP(7-39). (*A-B*)** Representative snapshots from MD simulations showing key peptide-binding residues close to the receptor core. TIP39 is depicted in green cartoon, TIP(7-39) is in wheat. Key residues are shown as sticks. Salt bridges and hydrogen bonds are showed as dash lines. **(*C*)** Frequency of an extracellular domain residue of PTH2R interacting with TIP39 (green) or TIP(7-39) (yellow) in simulations. The frequency value indicates the stability of a particular residue-peptide interaction. A large interacting frequency indicates a stable interaction.

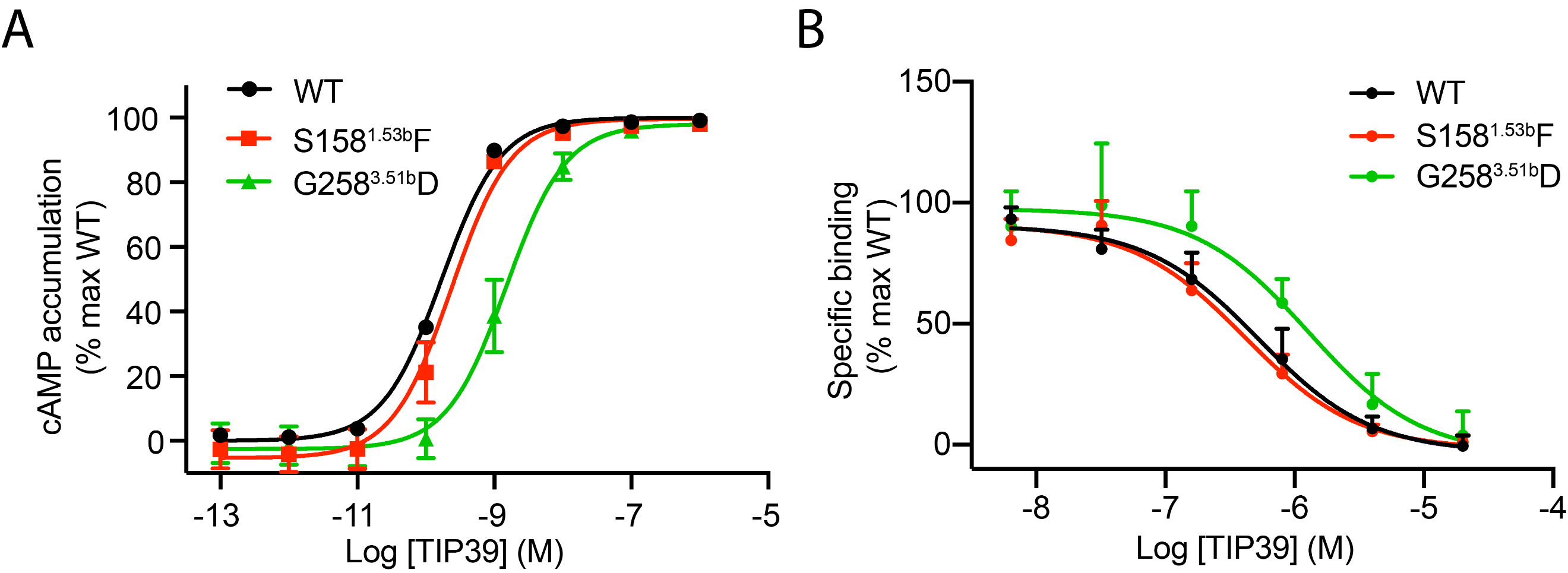

**Fig. S9. Effects of TIP39-mediated cAMP accumulation and binding affinity in disease-associated mutation. (*A*)** cAMP accumulation in wild-type (WT) and (single-point) mutant PTH2R expressing cells. **(*B*)** FITC-labelled ligand binding properties of WT and mutant PTH2Rs. Concentration-response curves were generated and graphed as means ± S.E.M. from three independent experiments.

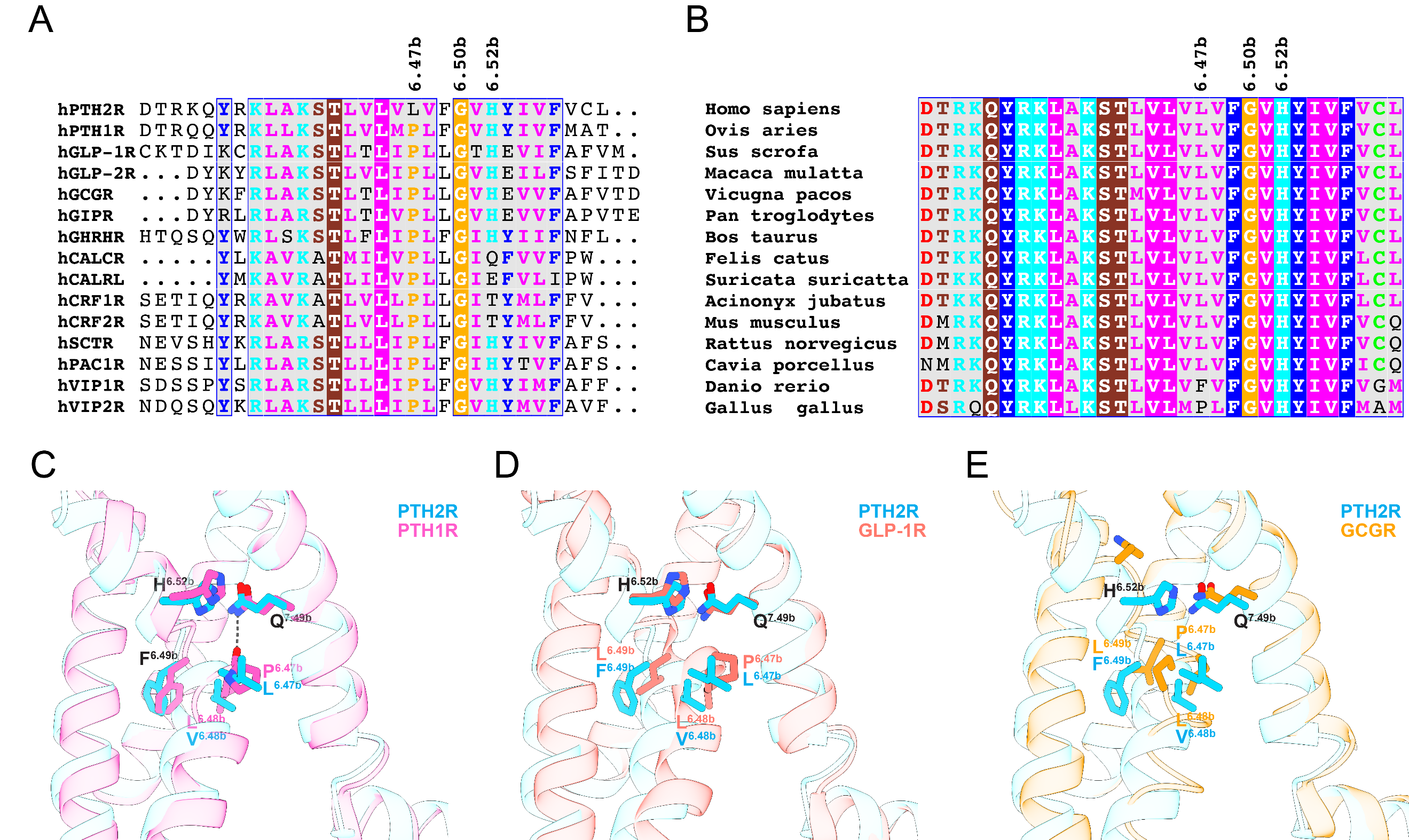

**Fig. S10. TM6 kink in class B1 GPCRs. (*A*)** Structure-based sequence alignment of TM6 in class B1 GPCRs. **(*B*)** Sequence conservation analysis: L^6.47b^XXG^6.50b^ motif in PTH2R across diversified species. **(*C-E*)**, Structural comparison of LXXG motif in PTH2R with PXXG motifs in PTH1R (*left*), GLP-1R (*middle*), and GCGR (*right*), respectively. ECD, TM1, TM4, agonist and G protein are omitted for clarity.

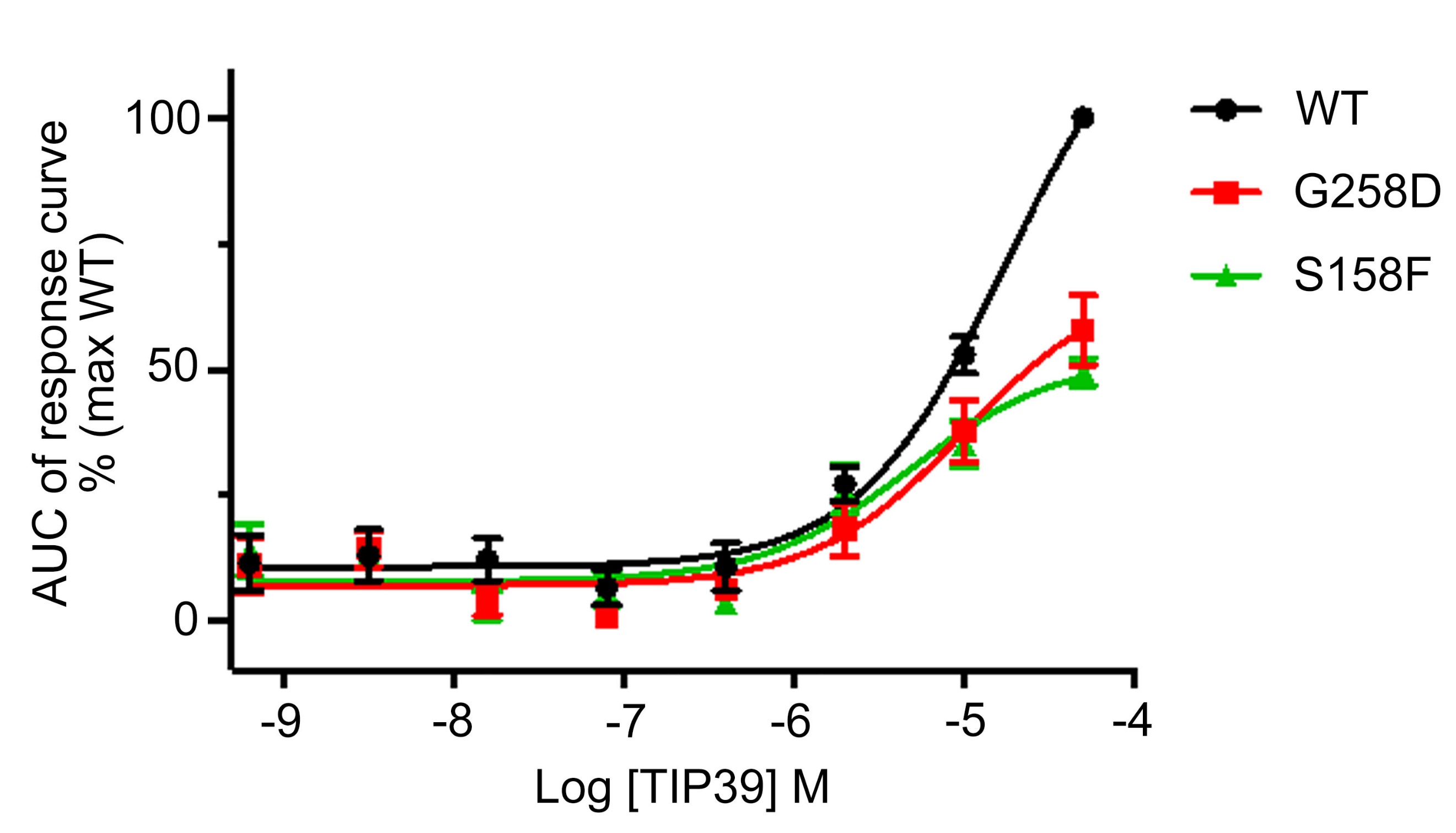

**Fig. S11. Effects of the disease-associated mutations on the TIP39-induced G_q_ protein coupling.** Concentration-response curves are expressed as AUC (area-under-the-curve) across the time-course response curve (0-13.5 min) for each concentration and normalized to the wild-type (WT) PTH2R.

**Table S1. Cryo-EM data collection, refinement and validation statistics.**

| **TIP39-PTH2R-G_s_ complex** | **Value** |
| --- | --- |
| **Data collection and processing** |  |
| Magnification | 95,694 |
| Voltage (kV) | 300 |
| Electron exposure (e^–^/Å^2^) | 80 |
| Defocus range (μm) | -1.2 ~ -2.2 |
| Pixel size (Å) | 0.5225 |
| Symmetry imposed | C1 |
| Initial particle projections (no.) | 3,241,855 |
| Final particle projections (no.) | 301,769 |
| Map resolution (Å) | 2.8 |
| FSC threshold | 0.143 |
| Map resolution range (Å) | 2.4-6.0 |
| **Refinement** |  |
| Initial model used (PDB code) | 6NBF |
| Model resolution (Å)  FSC threshold | 3.0  0.143 |
| Model resolution range (Å) | 3.0-5.0 |
| Map sharpening *B* factor (Å^2^) | -88.5 |
| Model composition  Non-hydrogen atoms  Protein residues  Lipids  Water | 9490  1160  0 |
| *B* factors (Å^2^)  Protein  Lipids | 98.0  95.9 |
| R.M.S. deviations  Bond lengths (Å)  Bond angles (°) | 0.005  1.119 |
| Validation  MolProbity score  Clash score  Poor rotamers (%) | 1.29  5.06  0.0 |
| Ramachandran plot  Favored (%)  Allowed (%)  Disallowed (%) | 97.9  2.1  0 |

**Table S2. Interaction between TIP39 and PTH2R.**

| **TIP39** | **PTH2R** | **Interaction** |
| --- | --- | --- |
| Ser1 | S310^ECL2^ | Side chain-side chain hydrogen bond |
|  | Q319^5.40b^ | Peptide amine terminus-side chain hydrogen bond |
| Ala3 | L323^5.44b^ | Hydrophobic interaction |
| Leu4 | L247^3.40b^ | Hydrophobic interaction |
|  | Y251^3.44b^ | Hydrophobic interaction |
|  | I322^5.43b^ | Hydrophobic interaction |
| Ala5 | E398^7.42b^ | Hydrophobic interaction |
| Asp6 | R190^2.60b^ | Salt bridge |
|  | Y152^1.47b^ | Side chain-side chain hydrogen bond |
| Asp7 | Y318^5.39b^ | Side chain-side chain hydrogen bond |
| Ala9 | M395^7.39b^ | Hydrophobic interaction |
|  | L399^7.43b^ | Hydrophobic interaction |
| Phe10 | K197^2.67b^ | Cation-pi stacking |
|  | F243^3.36b^ | Hydrophobic interaction |
| Arg11 | W307^45.51b^ | Side chain-backbone hydrogen bond |
| Glu12 | K137^1.32b^ | Side chain-side chain hydrogen bond |
| Arg13 | E142^1.37b^ | Salt bridge |
|  | Y145^1.40b^ | Cation-pi stacking |
| Leu16 | F141^1.36b^ | Hydrophobic interaction |
|  | K137^1.32b^ | Hydrophobic interaction |
| Glu21 | R305^ECL2^ | Salt bridge |
| Arg22 | D92^ECD^ | Side chain-side chain hydrogen bond |
| Arg23 | D92^ECD^ | Side chain-side chain hydrogen bond |
|  | Q130^1.25b^ | Side chain-side chain hydrogen bond |
| Trp25 | I34^ECD^ | Hydrophobic interaction |
|  | I38^ECD^ | Hydrophobic interaction |
| Leu26 | F93^ECD^ | Hydrophobic interaction |
| Met30 | L70^ECD^ | Hydrophobic interaction |
|  | F93^ECD^ | Hydrophobic interaction |
|  | Y122^ECD^ | Hydrophobic interaction |
|  | L126^ECD^ | Hydrophobic interaction |

**Table S3. Effect of PTH2R truncation on TIP39-mediated cAMP accumulation and ligand binding affinity^a^.**

|  | **cAMP accumulation** | | **Binding** |
| --- | --- | --- | --- |
|  | **pEC_50_ ± SEM** | **E_max_ (% WT)** | **pIC_50_ ± SEM** |
| PTH2R (25-550) (WT) | 9.2 ± 0.1 | 100 | 6.6 ± 0.2 |
| PTH2R (25-442)-20AA-LgBiT | 8.0 ± 0.1** | 109.8 ± 1.3** | 6.2 ± 0.4 |

^a^All data were fitted to a three-parameter logistic curve to obtain pEC_50_ and pIC_50_ values. Data represent means ± S.E.M. of at least three independent experiments performed duplicate. Statistical significance was determined with a two-tailed Student’s *t*-test. ***P* < 0.01. WT, wild-type.

**Table S4. Effects of residue mutation in the ligand-binding pocket on PTH2R-induced cAMP accumulation^a^.**

| **Receptor mutant** | **TIP39** | |
| --- | --- | --- |
|  | **pEC_50_±SEM** | **E_max_ (% WT)** |
| PTH2R | 9.2±0.08 | 100.2±0 |
| F141A | 7.7±0.1*** | 102.5±6.2 |
| E142A | 8.6±0.04 | 107.0±4.7 |
| Y152A | N.D. | N.D. |
| R190A | 6.3±0.05*** | 143.0±7.8** |
| K197A | 8.1±0.04*** | 94.8±0.8 |
| H202A | 8.6±0.2 | 100.7±8.1 |
| F243A | N.D. | N.D. |
| R305A | 8.0±0.01*** | 99.6±3.8 |
| L309A | 8.4±0.2* | 97.2±4.4 |
| Q319A | 8.8±0.4 | 100.1±1.6 |
| H384A | 8.6±0.2 | 104.0±4.4 |
| W391A | 8.3±0.06** | 105.0±3.7 |
| M395A | 7.6±0.06*** | 105.6±7.1 |
| L399A | 7.3±0.02*** | 105.6±3.6 |

^a^All data were fitted to a three-parameter logistic curve to obtain pEC_50_ values. Data represent means ± S.E.M. of at least three independent experiments performed duplicate. One-way ANOVA and Dunnett’s post-test were used to determine statistical difference. **P*<0.05, ***P*<0.01, ****P*<0.001. WT, wild-type; N.D., values that could not be determined due to incomplete curve fitting.

**Table S5. Effects of residue mutation on ligand recognition and receptor specificity^a^.**

| **Receptor mutant** | **TIP39** | | **PTH(1-34)** | |
| --- | --- | --- | --- | --- |
|  | **pEC_50_±SEM** | **E_max_ (% WT)** | **pEC_50_±SEM** | **E_max_ (% WT)** |
| PTH2R | 10.4±0.07 | 100 | 9.1±0.12 | 100 |
| I34A | 9.8±0.20 | 110.8±0.78 | 8.4±0.18 | 111.3±2.16 |
| I38A | 8.8±0.13* | 103.2±3. | 7.9±0.10** | 108.3±5.43 |
| Y152A | N.D. | N.D. | N.D. | N.D. |
| R190A | 7.1±0.34*** | 108.5±5.89 | 6.8±0.04*** | 106.2±4.56 |
| K197A | 8.8±0.10* | 97.6±1.16 | 8.2±0.51* | 98.2±3.80 |
| R305A | 9.1±0.04 | 94.6±1.38 | 9.1±0.08 | 98.1±2.20 |
| Y318A | 7.5±0.12*** | 100.3±1.19 | 9.7±0.10 | 101.6±2.10 |
| L399A | 8.2±0.04** | 93.0±5.63 | N.D. | N.D. |

^a^All data were fitted to a three-parameter logistic curve to obtain pEC_50_ values. Data represent means ± S.E.M. of at least three independent experiments performed duplicate. One-way ANOVA and Dunnett’s post-test were used to determine statistical difference. **P*<0.05, ***P*<0.01, ****P*<0.001. WT, wild-type; N.D., values that could not be determined due to incomplete curve fitting.

**Table S6. Effects of disease-associated mutation on TIP39-mediated cAMP accumulation, ligand binding affinity and G_s_ coupling^a^.**

| **Receptor mutant** | **cAMP accumulation** | **Binding** | **G_s_ coupling** | |
| --- | --- | --- | --- | --- |
|  | **pEC_50_±S.E.M.** | **pIC_50_±S.E.M.** | | **pEC_50_±S.E.M.** |
| PTH2R | 9.8±0.1 | 6.4±0.2 | | 5.8±0.3 |
| S158F | 9.7±0.1 | 6.4±0.2 | | 5.2±0.1 (P=0.2302) |
| G258D | 8.8±0.2 (P=0.0040) | 5.9±0.1 (P=0.2726) | | 5.1±0.3 (P=0.1426) |

^a^All data were fitted to a three-parameter logistic curve to obtain pEC_50_ and pIC_50_ values. Data represent means ± S.E.M. of at least three independent experiments performed duplicate. One-way ANOVA and Dunnett’s post-test were used to determine statistical difference. WT, wild-type.

**Supplementary Materials and Methods**

**Cell Line.** High Five insect cells (ThermoFisher Scientific) were grown in ESF 921 serum-free media (Expression Systems) at 27°C and 120 rpm. Functional studies were performed in HEK-293T cells purchased from the Cell Bank at the Chinese Academy of Sciences (CAS; Shanghai). The cells were cultured in Dulbecco’s modified Eagle’s medium (DMEM; Life Technologies) supplemented with 10% (v/v) fetal bovine serum (Gibco) at 37°C and 5% CO_2_ in a humidified incubator.

**Receptor expression.** PTH2R and G_s_ heterotrimer were co-expressed in High Five insect cells. We prepared the viruses using Bac-to-Bac system (Invitrogen). The cells were grown to a density of 2.5 × 10^6^ -3.0 × 10^6^ cells/mL and infected with four types of baculoviruses at a ratio of 1:2:2:2 for PTH2R, DNGαs, Gβ1 and Gγ2, respectively. The cells were collected by centrifugation after 48 h and cell pellets were stored at -80°C.

**Production of Nb35.** Nb35 was expressed in *E. coli* strain BL21 and purified by nickel chromatography according to previously described methods (1). Crude sample was subjected to size-exclusion chromatography HiLoad 16/600 Superdex 200 pg column (GE healthcare) with running buffer of 20 mM HEPES, pH 7.4, and 100 mM NaCl. Selected fractions of Nb35 were concentrated to 1.2 mg/mL supplemented with 30% (v/v) glycerol and stored at -80°C after flash frozen. The quality of purified proteins was assessed by SDS-PAGE.

**Image processing.** Dose-fractionated image stacks were subjected to beam-induced motion correction using MotionCor2.1 (2). A sum of all frames, filtered according to the exposure dose, in each image stack was used for further processing. Contrast transfer function parameters for each micrograph were determined by Gctf v1.06 (3). Particle selection, 2D and 3D classifications were performed on a binned dataset with a pixel size of 2.09 Å using RELION-3.0-beta2 (4). Auto-picking yielded 3,241,855 particle projections were subjected to reference-free 2D classification to discard false positive particles or particles categorized in poorly defined classes, producing 2,395,321 particle projections for further processing. This subset of particle projections was subjected to a round of maximum-likelihood-based three dimensional classifications with a pixel size of 2.09 Å, resulting in one well-defined subsets with 879,801 projections. Further 3D classifications with mask on the complex produced one good subset accounting for 575,126 particles, which were subsequently subjected to a round of 3D classifications with mask on the receptor. A selected subset containing 301,769 projections was then subjected to 3D refinement and Bayesian polishing with a pixel size of 1.045 Å. After last round of refinement, the final map has an indicated global resolution of 2.82 Å at a Fourier shell correlation (FSC) of 0.143. Local resolution was determined using the Bsoft package with half maps as input maps (5).

**Molecular dynamics simulation and analysis.** On the basis of the CHARMM36m all-atom force field (6-8), molecular dynamics simulations were conducted using GROMACS 5.1.4 (9, 10). After 100 ns equilibration, a 1 μs production run was carried out for each simulation. All productions were carried out in the NPT ensemble at temperature of 303.15 K and a pressure of 1 atm. Temperature and pressure were controlled using the velocity-rescale thermostat (11) and the Parrinello-Rahman barostat with isotropic coupling (12), respectively. Equations of motion were integrated with a 2 fs time step, the LINCS algorithm was used to constrain bond length (13). Non-bonded pair lists were generated every 10 steps using distance cut-off of 1.4 nm. A cut-off of 1.2 nm was used for Lennard-Jones (excluding scales 1-4) interactions, which were smoothly switched off between 1 and 1.2 nm. Electrostatic interactions were computed using particle-mesh-Ewald algorithm with a real-space cutoff of 1.2 nm. The last 200 ns trajectory of each simulation was used to calculate average values. The non-hydrogen atom distance between two residues is the minimal distance between non-hydrogen atoms from two different residues; the distance between the non-hydrogen atoms from the same residue is excluded. The backbone distance between two motifs is the minimal distance between backbone atoms from two different defined motifs; the distance between the backbone atoms from the same motif is excluded. The frequency of a particular residue interacting with a peptide was calculated by counting how many times this residue interacts with the peptide in the simulation snapshots. The interaction is defined by the heavy atom distance between the residue and the peptide using 4 Å as cutoff. The interacting frequency value indicates the stability of a particular residue-peptide interaction. A large interacting frequency indicates a stable interaction.

**SI References**

1.	E. Pardon *et al.*, A general protocol for the generation of nanobodies for structural biology. *Nat Protoc* **9**, 674-693 (2014).

2.	S. Q. Zheng *et al.*, MotionCor2: anisotropic correction of beam-induced motion for improved cryo-electron microscopy. *Nat Methods* **14**, 331-332 (2017).

3.	K. Zhang, Gctf: Real-time CTF determination and correction. *J Struct Biol* **193**, 1-12 (2016).

4.	S. H. Scheres, RELION: implementation of a bayesian approach to cryo-EM structure determination. *J Struct Biol* **180**, 519-530 (2012).

5.	J. B. Heymann, Guidelines for using Bsoft for high resolution reconstruction and validation of biomolecular structures from electron micrographs. *Protein Sci* **27**, 159-171 (2018).

6.	O. Guvench *et al.*, CHARMM additive all-atom force field for carbohydrate derivatives and its utility in polysaccharide and carbohydrate-protein modeling. *J Chem Theory Comput* **7**, 3162-3180 (2011).

7.	J. Huang *et al.*, CHARMM36m: an improved force field for folded and intrinsically disordered proteins. *Nat Methods* **14**, 71-73 (2017).

8.	A. D. MacKerell *et al.*, All-atom empirical potential for molecular modeling and dynamics studies of proteins. *J Phys Chem B* **102**, 3586-3616 (1998).

9.	D. Van Der Spoel *et al.*, GROMACS: fast, flexible, and free. *J Comput Chem* **26**, 1701-1718 (2005).

10.	B. Hess, C. Kutzner, D. van der Spoel, E. Lindahl, GROMACS 4: Algorithms for highly efficient, load-balanced, and scalable molecular simulation. *Journal of Chemical Theory and Computation* **4**, 435-447 (2008).

11.	G. Bussi, D. Donadio, M. Parrinello, Canonical sampling through velocity rescaling. *J Chem Phys* **126** (2007).

12.	K. M. Aoki, F. Yonezawa, Constant-pressure molecular-dynamics simulations of the crystal-smectic transition in systems of soft parallel spherocylinders. *Phys Rev A* **46**, 6541-6549 (1992).

13.	B. Hess, P-LINCS: A parallel linear constraint solver for molecular simulation. *J Chem Theory Comput* **4**, 116-122 (2008).
